# Dynamic allocation of DnaK couples proteostasis demand to chaperone availability

**DOI:** 10.64898/2026.07.01.735817

**Authors:** Ada Pajari, Taras Redchuk, Jarno Mäkelä

**Affiliations:** Department of Neuroscience and Biomedical Engineering, Aalto University, Espoo, Finland; Institute of Biotechnology, Helsinki Institute of Life Sciences, University of Helsinki, Helsinki, Finland

## Abstract

Protein-quality-control systems are essential for cells to maintain protein homeostasis during both steady-state growth and acute stress. Yet, how protein chaperone engagement dynamically changes as proteostasis demand increases remains poorly understood. Here, we combine rapid temperature control with single-molecule tracking to follow the bacterial heat shock protein 70 (Hsp70) DnaK in live *Escherichia coli*. We found that most DnaK molecules actively engage with the proteome already at the optimal growth temperature, rather than form a freely diffusing reserve. Acute heat shock reallocates the DnaK pool within seconds primarily increasing the lifetime but also frequency of client engagements. Perturbing the chaperone network reveals that this redistribution reflects proteostasis demand and network capacity: loss of small heat shock proteins IbpAB drives prolonged DnaK engagement and limits recovery, whereas overexpression of thermolabile proteins alone can produce heat shock-like dynamics. Together, our findings reveal how chaperones are dynamically reallocated during proteotoxic stress and establish DnaK mobility as a sensitive readout of proteostasis demand.

## INTRODUCTION

Protein homeostasis, or proteostasis, is essential for all living cells to ensure that proteins fold, function, and remain soluble in a crowded intracellular environment. This challenge becomes acute during proteotoxic stress, when changes in temperature, metabolism, or protein expression can rapidly increase the number of partially unfolded and aggregation-prone proteins^1,2^. Cells must therefore not only regulate the abundance of chaperones, but also dynamically allocate them among competing protein-folding demands. How this allocation occurs in living cells, and how rapidly it can be remodeled as proteostasis demand changes, remains poorly understood.

A central component of the bacterial proteostasis network is the heat-shock protein 70 (Hsp70) chaperone DnaK. DnaK is an abundant ATP-dependent chaperone that recognizes hydrophobic regions exposed during protein synthesis, unfolding or misfolding. Client binding is coupled to an ATPase cycle regulated by the J-domain co-chaperone DnaJ and the nucleotide-exchange factor GrpE^3–6^. Proteome-scale studies in *Escherichia coli* have identified hundreds of DnaK-interacting proteins, including aggregation-prone clients that depend strongly on DnaK for folding and maintenance^7,8^. DnaK also cooperates with the small heat-shock proteins IbpA and IbpB, the GroEL/ES complex, the disaggregase ClpB and the bacterial Hsp90 homologue HtpG, and regulates the heat-shock sigma factor σ^32^, linking the folding state of the proteome to transcriptional induction of the heat-shock response^9–16^.

Despite extensive study of the roles of DnaK in proteostasis, it is unknown how DnaK distributes between its activities in the cell during balanced growth and how this is remodeled during acute proteotoxic stress. Much of our understanding of Hsp70/DnaK mechanisms derives from *in vitro* studies that lack the cellular context, the complexity of the proteome, and the presence of many co-chaperones. Proteostasis demand should alter not only how much DnaK is engaged, but also how long individual DnaK molecules dwell in distinct states, how these states interconvert, and where these states are enriched within the cell. Recent live-cell single-molecule approaches have begun to reveal the interaction dynamics of individual chaperone systems. Single-molecule tracking of bacterial Trigger Factor revealed rapid exchange with ribosomes and nascent polypeptides in *E. coli*^17^, whereas studies of the eukaryotic TRiC chaperonin showed repeated, short-lived probing interactions whose lifetimes increased for folding-challenged clients^18^.

Here, we combine rapid temperature control with live-cell single-molecule tracking to determine how DnaK interaction dynamics change as proteostasis demand changes in *E. coli* (**Figure 1A and B**). We show that DnaK activity in cells is not restricted to stress conditions; DnaK is already broadly engaged with the cellular proteome during optimal growth. Heat shock rapidly remodels this basal interaction landscape by shifting DnaK towards a slow population with increased dwell times and peripheral localization. We further perturb the proteostasis network in two complementary ways: by reducing chaperone-network capacity through removal of IbpAB, DnaJ, or HtpG, and by increasing folding demand through overexpression of thermolabile proteins. Together, these findings reveal how bacteria dynamically allocate their chaperone pools during steady-state growth and proteotoxic stress, and show that DnaK mobility provides a sensitive readout of proteostasis demand.

**Figure 1.**
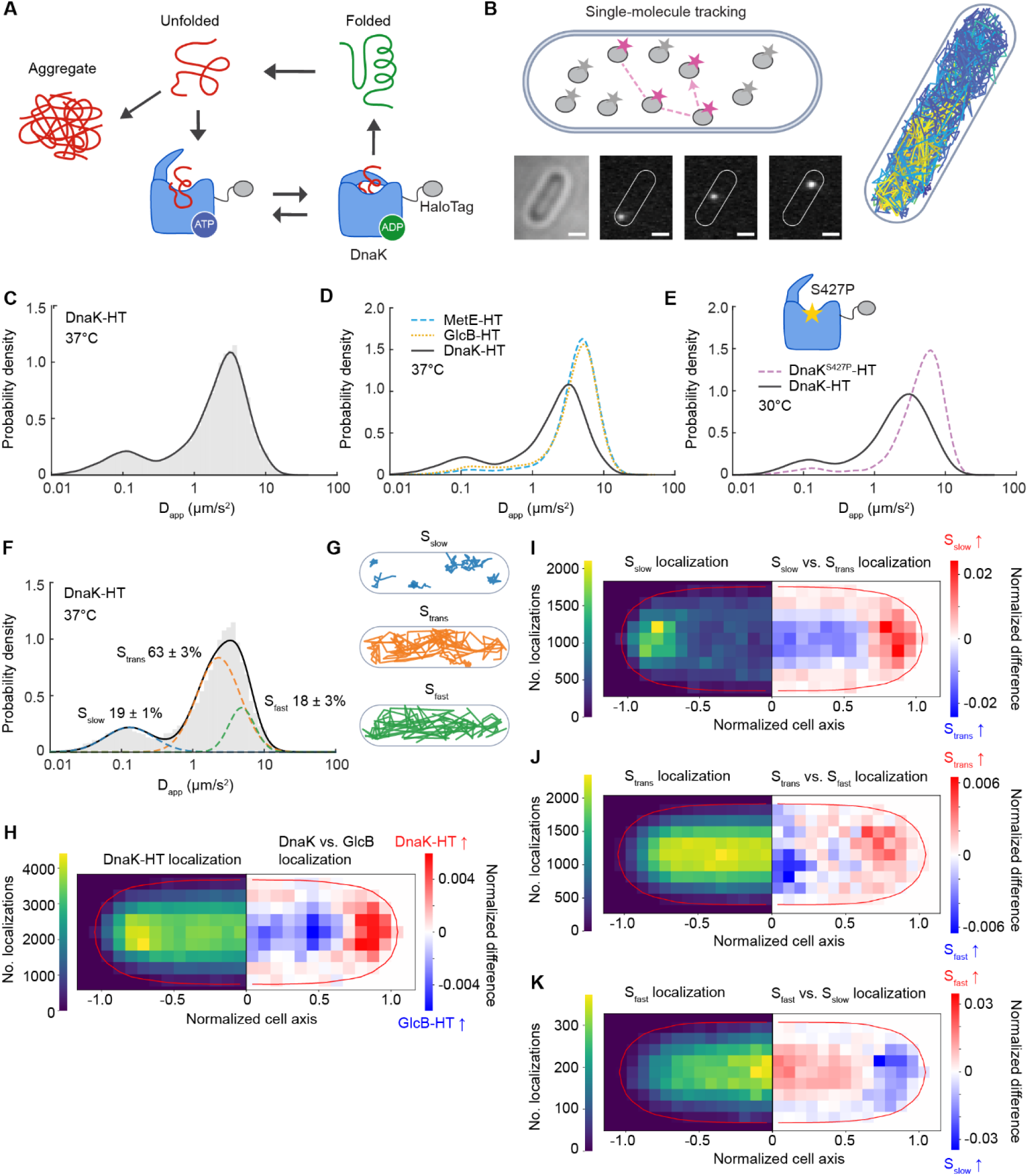
Single-molecule dynamics of DnaK. (**A**) Illustration of DnaK chaperone function. (**B**) Schematic of single-molecule tracking (scale bars 1 µm). (**C**) Apparent diffusion coefficient (D_app_) distribution (log_10_ scale) of JFX554-labeled DnaK-HaloTag at 37°C (25324 tracks in 562 cells, JML_Ec_05). Mean D_app_: 2.76 ± 0.05 µm^2^/s. (**D**) D_app_ of GlcB-HaloTag (48682 tracks in 891 cells, JML_Ec_141) and MetE-HaloTag (43634 tracks in 1052 cells, JML_Ec_136) at 37°C. Mean D_app_: 4.64 ± 0.03 µm^2^/s and 4.89 ± 0.13 µm^2^/s, respectively. Also shown is the DnaK-HaloTag distribution from (C). (**E**) D_app_ of substrate-binding-deficient DnaK^S427P^-HaloTag (18590 tracks in 944 cells, JML_Ec_164) at 30°C. Mean D_app_: 5.24 ± 0.37 µm^2^/s. Also shown wild-type DnaK-HaloTag at 30°C. (**F**) D_app_ of DnaK-HaloTag at 37°C fitted by a three-state Gaussian mixture model (GMM). The fitted mean D_app_ for the slow (S_slow_), transient (S_trans_), and fast diffusion states (S_fast_) were 0.13 ± 0.01 µm^2^/s; 2.29 ± 0.12 µm^2^/s; 4.83 ± 0.06 µm^2^/s, respectively. (**G**) Example single-molecule trajectories from each diffusional state of DnaK (< 0.45 µm^2^/s, 0.45 < D_app_ < 5.4 µm^2^/s, and > 5.4 µm^2^/s). (**H**) DnaK-HaloTag localization at 37 °C (left side) and normalized difference between DnaK-HaloTag and GlcB-HaloTag distributions (right side). Red indicates higher DnaK occupancy while blue indicates higher GlcB occupancy. (**I-K**) Localizations of diffusional states S_slow_, S_trans_ and S_fast_ (left side) and normalized differences between the states (right side) at 37°C. Localizations are plotted on normalized cell coordinates. All data are from three biological replicates and ± indicates the standard error of the mean [SEM] between the replicates.

## RESULTS

### DnaK samples the proteome during steady-state growth

To explore DnaK dynamics, we performed single-molecule tracking in live *E. coli* cells grown in M9 minimal medium supplemented with glycerol, casamino acids and thiamine (M9glyCAAT) at 37°C. We introduced a HaloTag fusion to the C-terminus of DnaK through genetic modification at the native chromosomal locus. The resulting DnaK-HaloTag fusion did not affect cell survival during a semi-lethal heat shock (15 min at 51°C) in comparison to wildtype (**Figure 1 – figure supplement 1**), indicating preserved DnaK functionality in the fusion. For imaging, DnaK-HaloTag was fluorescently labelled using the membrane-permeable Janelia Fluor 554 dye (JFX554)^19^, and the local temperature at the cells was controlled by the VAHEAT system with an optically transparent thin-film heater and integrated temperature sensor at the coverslip.

Based on the known substrate-binding cycle of DnaK^3^, we expected at least two mobility populations: a slow state representing substrate-DnaK interactions and a faster diffusive state. Single-molecule tracking revealed a mean apparent diffusion coefficient of *D_app_* = 2.76 ± 0.05 µm^2^/s (±SEM), with a small low mobility peak and a larger fast population (**Figure 1C**). However, even molecules in the fast DnaK population (∼3.7 µm^2^/s) diffused more slowly than expected for a freely diffusing protein of comparable size^20–22^, suggesting that rapid DnaK-client interactions may also contribute to this population.

To distinguish interaction-dependent slowing from effects of molecular size, we compared DnaK with similarly sized HaloTag fusions of GlcB (80 kDa) and MetE (85 kDa), proteins expected to diffuse largely freely in the cytoplasm^20^. Consistent with this, both showed a single fast-moving population with similar diffusion coefficients (MetE: *D_app_* = 4.89 ± 0.13 µm^2^/s; GlcB: *D_app_* = 4.64 ± 0.03 µm^2^/s) (**Figure 1D, Figure 1 - figure supplement 2A-B**). Despite their comparable size, both diffused faster than DnaK, supporting the conclusion that DnaK mobility is reduced by frequent interactions with cellular clients.

To further test whether reduced DnaK mobility reflects substrate binding, we examined DnaK^S427P^, a substrate-binding-domain mutant with reduced client affinity but near-wild-type ATPase activity^23,24^. Because DnaK^S427P^ cells became filamentous at higher temperatures, DnaK^S427P^ and wild-type DnaK were compared at 30°C. DnaK^S427P^ was substantially more mobile than wild type (*D_app_* = 5.24 ± 0.37 µm^2^/s versus 3.01 ± 0.08 µm^2^/s) (**Figure 1E, Figure 1 - figure supplement 2C-D**), supporting the idea that rapid client interactions contribute to the reduced mobility of DnaK. A second substrate-binding-deficient mutant, DnaK^V436F^,^25^ showed only a modest increase in mobility (3.37 ± 0.10 µm^2^/s, **Figure 1 – figure supplement 2E**), indicating a weaker effect on DnaK-client interactions *in vivo*.

We next quantified the occupancy of distinct DnaK mobility states using a three-component Gaussian mixture model (GMM) (**Figure 1F**). The high-mobility component was constrained using GlcB as a size-matched freely diffusing reference, with MetE showing comparable diffusion (**Figure 1 – figure supplement 2A, B, F**). The model resolved slow (S_slow_, 19 ± 1%), intermediate (S_trans_, 63 ± 3%), and fast (S_fast_, 18 ± 3%) populations (**Figure 1G**). S_slow_, which was nearly abolished in DnaK^S427P^, is consistent with prolonged engagement with clients or larger proteostasis assemblies, whereas S_trans_ is consistent with short-lived interactions and S_fast_ predominantly with free diffusion. Thus, most DnaK molecules actively engage with clients at any given time even at the optimal growth temperature.

Although bacteria lack membrane-bound organelles, damaged proteins and aggregates can become spatially organized, particularly at the cell poles or in nucleoid-free regions^26–28^. We therefore asked whether DnaK mobility states occupy distinct intracellular regions. Unlike the freely diffusing proteins GlcB and MetE, DnaK was enriched toward the cell poles and periphery already at 37°C (**Figure 1H, Figure 1 - figure supplement 3A-B**). The individual mobility states showed distinct spatial patterns (**Figure 1I-K**): S_trans_ was more polar and peripheral than S_fast_, whereas S_slow_ showed the strongest polar enrichment. Thus, even under balanced growth, DnaK is spatially partitioned among mobility states consistent with different modes of proteome engagement. We next asked how these interactions change when proteostasis demand increases abruptly.

### Heat shock rapidly redistributes the DnaK pool

Given its central role in the heat shock response, we next examined how DnaK dynamics change following an acute, non-lethal temperature upshift from 37°C to 45°C. Cells were heated to 45°C on the microscope with an average ramp time of 7.0 ± 1.2 s (±SEM), and single-molecule tracking data were collected before and after the upshift (**Figure 2A**). DnaK mobility decreased immediately upon reaching 45°C, with the mean D_app_ falling from 2.76 ± 0.05 µm^2^/s to 1.11 ± 0.33 µm^2^/s, consistent with the increased engagement caused by heat-induced proteostasis stress. This agrees with biochemical studies showing that elevated temperature shifts the DnaK/DnaJ/GrpE system toward a high-affinity substrate-sequestering state^29^.

**Figure 2.**
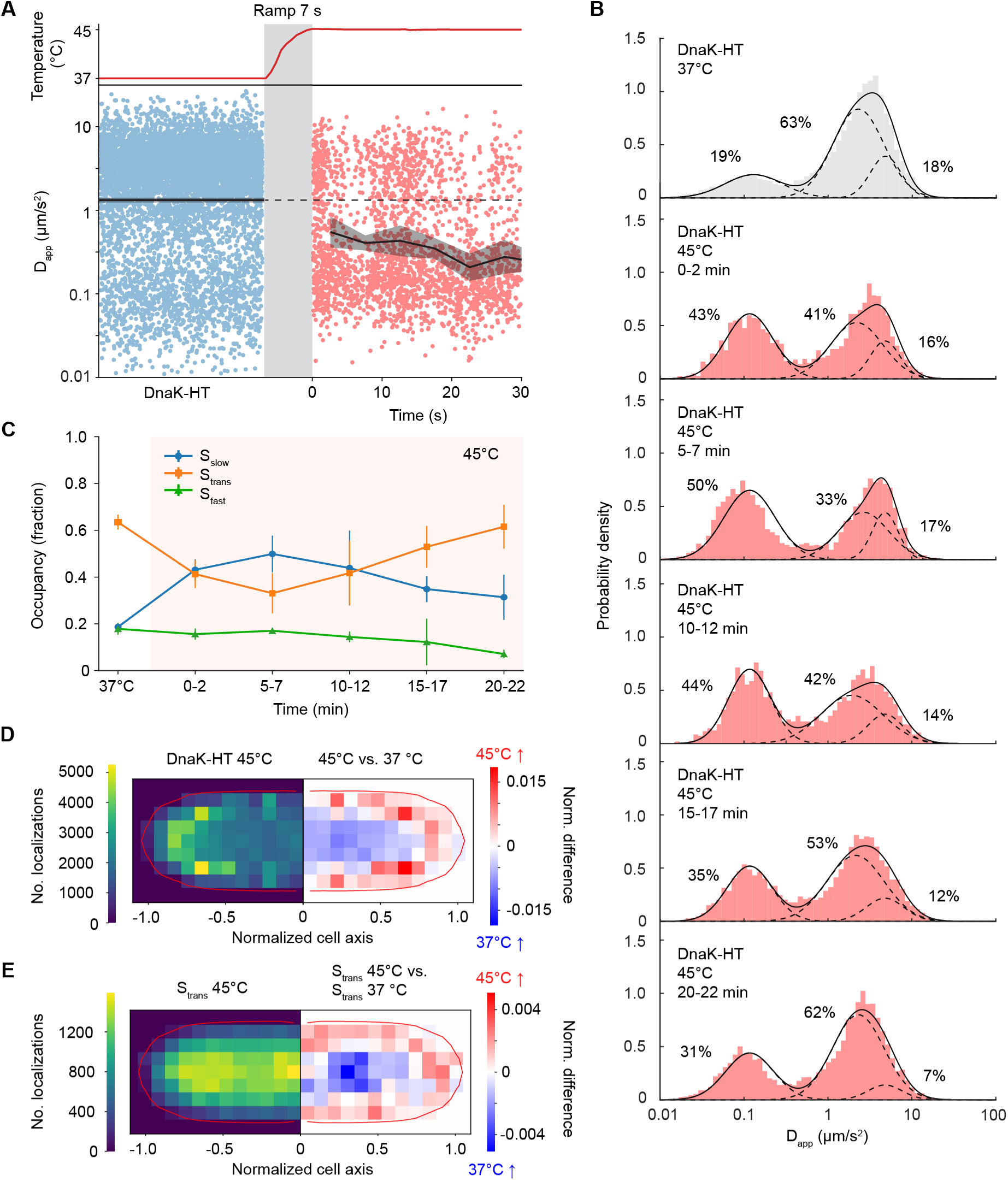
Diffusional dynamics of DnaK-HaloTag during heat shock. **(A)** Top: Temperature read-out during upshift from 37°C to 45°C. Bottom: DnaK-HaloTag single-molecule tracks at 37°C (blue) and at the onset of 45°C heat shock (red). The black line denotes the mean and the shaded areas denote the standard error of the mean (SEM). **(B)** D_app_ distributions fitted with a three-state GMM at 37°C and during a 45°C heat shock time course (18962 tracks and 869 cells). Data statistics are available in Supplementary Table 1-2. **(C)** GMM-derived occupancy for each diffusional state of DnaK-HaloTag at 37°C and during the 45°C heat shock. Error bars indicate the standard error of the mean [SEM] between the replicates. **(D)** Localizations of DnaK-HaloTag at 45°C (left side) and normalized difference between 45°C and 37°C (right side). Red indicates higher DnaK occupancy at 45°C while blue indicates higher DnaK occupancy at 37°C. (**E**) Localizations of S_trans_ at 45°C (left side) and normalized difference between 45°C and 37°C (right side). Red indicates higher occupancy at 45°C while blue indicates higher occupancy at 37°C. Localizations are plotted on normalized cell coordinates. All data are from three biological replicates.

We then followed DnaK dynamics over 20 min at 45°C, sampling every 5 min (**Figure 2B**). Three-component GMM analysis showed a rapid increase in S_slow_ occupancy, which peaked after 5-7 min, and then partially declined but remained above the 37°C baseline (**Figure 2B, C**). S_trans_ showed the opposite trend, decreasing during the first 5-7 min before recovering by 20 min, whereas S_fast_ gradually decreased throughout the time course. Thus, DnaK did not fully return to its pre-stress state but instead adapted to a new state characterized by enrichment of the low-mobility population and depletion of the fast population.

Heat shock also altered DnaK localization. DnaK became more enriched at the cell poles and periphery (**Figure 2D**), reflecting both increased occupancy of the S_slow_ state and a more peripheral distribution of the S_trans_ population (**Figure 2E**, **Figure 2 – figure supplement 1A-B**). These results show that acute heat stress rapidly reorganizes both the dynamics and localization of DnaK, reallocating the pre-existing chaperone pool from proteome sampling toward more persistent client engagement.

### Heat shock extends DnaK interaction lifetimes

We next asked how individual DnaK molecules exchange between mobility states over time. State occupancy describes how much of the DnaK pool is committed to a given dynamic state at any moment, but does not reveal how rapidly those molecules become available again. Short-lived engagements allow DnaK to cycle rapidly between clients, while prolonged binding keeps individual molecules committed for longer and reduces their immediate availability for additional folding demands.

To quantify apparent dwell times and transition rates, we used the ExTrack Hidden Markov Model (HMM)^30^, which infers molecular state sequences from single-particle trajectories and accommodates noisy tracking data, heterogeneous diffusion within a state, and rapid transitions. Because HMM-based approaches can overestimate the number of states in noisy trajectories^31–33^, we used a coarse-grained three-state model corresponding to the slow, transient, and fast mobility states defined above.

At 37°C, all three states were short-lived, with apparent dwell times of 0.20 ± 0.02 s for S_trans_, 0.27 ± 0.07 s for S_fast_, and 0.20 ± 0.05 s for S_slow_ (**Figure 3A, Supplementary Table 3**). Transitions between S_fast_ and S_slow_ occurred predominantly through S_trans_, suggesting that the intermediate state forms a kinetic bridge between highly mobile DnaK and longer-lived client engagements. Together with the partially overlapping S_trans_ and S_fast_ diffusion distributions (**Figure 1F**), these rapid transitions are consistent with continuous proteome sampling through repeated client binding and release during normal growth.

**Figure 3.**
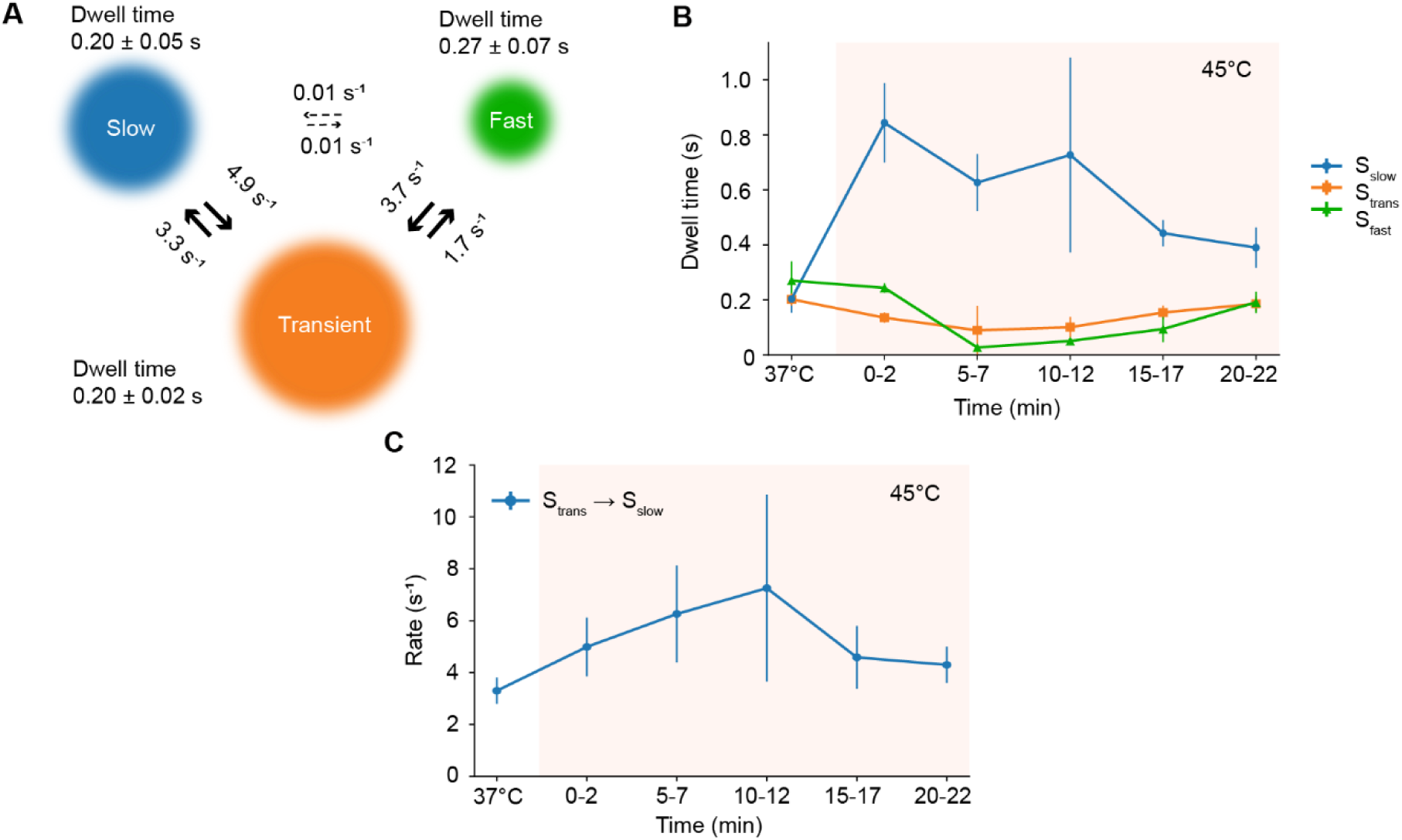
Hidden Markov Model analysis of DnaK state-transition dynamics. (**A**) State transition rates and dwell times obtained from a three-state Hidden Markov model analysis of DnaK data at 37°C. (**B**) Apparent dwell times of DnaK states at 37°C and during 45°C heat shock. (**C**) DnaK transition rate from S_trans_ into S_slow_ at 37°C and during 45°C heat shock. Error bars and ± indicate the standard error of the mean (SEM). Same data as in Figure 2. Data statistics are available in Supplementary Table 3-4.

Heat shock mostly affected S_slow_, whose dwell time increased 4.1-fold to 0.84 ± 0.14 s within the first 2 min after the temperature upshift (**Figure 3B**). Although it subsequently decreased over the rest of the time course, S_slow_ dwell time remained above the pre-stress level after 20 min at 45°C. In contrast, S_trans_ and S_fast_ dwell times decreased during the early heat-shock response, reached minima after 5–7 min and returned close to baseline by 20 min. Consistent with these longer-lived engagements, slow-moving DnaK also became more spatially concentrated during heat shock (**Figure 3 – figure supplement 1**). At 37 °C, S_slow_ and S_trans_ showed similar relative-density distributions, whereas heat shock produced a high-density tail specifically for S_slow_, indicating increased clustering of low-mobility DnaK.

Increased S_slow_ occupancy could arise either from longer residence in the state or from more frequent entry into the state. We therefore quantified S_trans_ → S_slow_ transition rates during the heat shock (**Figure 3C**). Entry rates increased more gradually than dwell time, reaching ∼2-fold maximum after 10-12 min of heat shock. This indicates that the rapid early increase in S_slow_ is driven mainly by prolonged engagement rather than by more frequent entry. Together with the state occupancy changes, these results indicate that heat shock reallocates the pre-existing DnaK pool by extending the lifetime of engagements, therefore reducing the pool available for rapid proteome sampling.

### IbpAB buffer aggregation-prone clients to preserve DnaK availability

The small heat-shock proteins IbpA and IbpB bind aggregation-prone substrates and have been proposed to cooperate with the DnaK system^14^. Given the increased spatial concentration of slow-moving DnaK during heat shock, we asked how IbpA behaves dynamically in living cells and how its dynamics relate to DnaK. We generated a functional endogenous IbpA-HaloTag fusion (**Figure 4 – figure supplement 1**) and performed single-molecule tracking at 37°C and during 45°C heat shock.

At 37°C, IbpA occupied three mobility states that were more clearly separated than those of DnaK, likely reflecting the smaller molecular mass and faster free diffusion of IbpA (**Figure 4A, B**). HMM analysis yielded apparent dwell times of 0.17 ± 0.03 s, 0.15 ± 0.01 s, and 0.027 ± 0.002 s for S_slow_, S_trans_, and S_fast_, respectively (**Figure 4C**). Approximately 80% of IbpA already occupied the slow or transient populations at 37°C. Together with the similar behavior of DnaK, this suggests that rapid transient probing may be a shared feature of chaperone dynamics.

**Figure 4.**
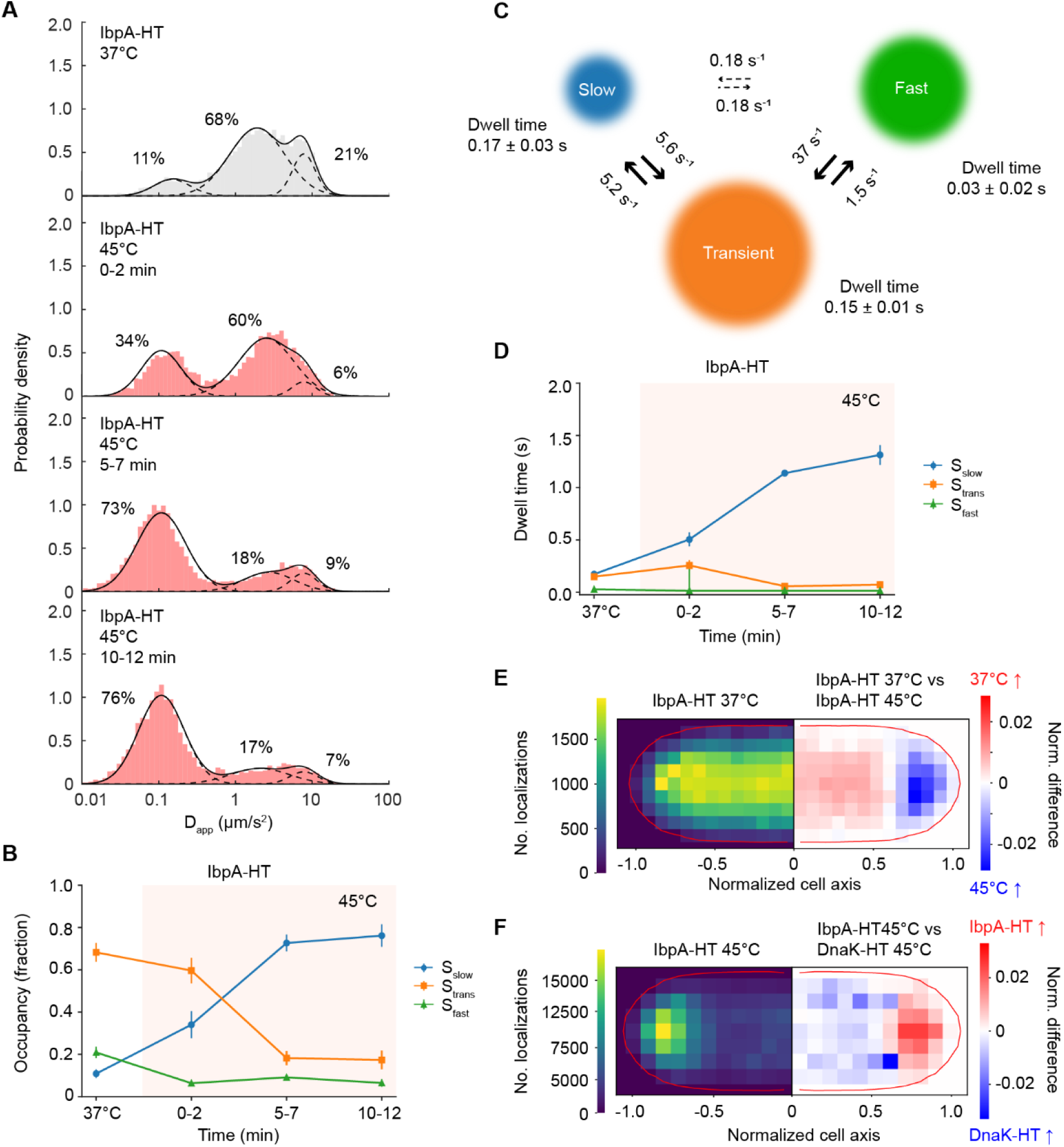
IbpA dynamics during steady-state growth and heat shock. (**A**) D_app_ distributions of IbpA-HaloTag (JML_Ec_14) at 37°C (26619 tracks in 303 cells) and during 45°C heat shock (16669 tracks in 880 cells). The D_app_ distributions were fitted by a three-state GMM. (**B**) GMM state occupancy for each IbpA-HaloTag diffusional state. (**C**) IbpA-HaloTag state transition rates obtained from a three-state Hidden Markov Model (HMM)-analysis of IbpA single-molecule data at 37°C. (**D**) HMM-derived dwell times of diffusional states of IbpA-HaloTag at 37°C and during 45°C heat shock. (**E**) IbpA-HaloTag localizations at 37°C (left side) and the normalized differences between 37 °C and 45°C (right side). Red indicates higher IbpA occupancy at 37°C while blue indicates higher IbpA occupancy at 45°C. (**F**) IbpA-HaloTag localizations at 45 °C (left side) and the normalized differences from DnaK-HaloTag localizations at 45°C (right side). Red indicates higher IbpA occupancy while blue indicates higher DnaK occupancy. Localizations are plotted on normalized cell coordinates. Data statistics are available in Supplementary Table 1-4. All data are from three biological replicates. Error bars and ± indicate the standard error of the mean [SEM] between the replicates.

IbpA responded more strongly and persistently to heat shock than DnaK. S_slow_ increased to ∼76% occupancy after 10–12 min at 45°C (**Figure 4B**) and encompassed almost all IbpA molecules after 15 min (**Figure 4 – figure supplement 2**). S_slow_ dwell time increased to 1.3 ± 0.1 s after 10–12 min and showed no recovery toward the pre-stress level (**Figure 4D**). This transition occurred more gradually than the rapid increase in DnaK S_slow_ occupancy, consistent with distinct roles of the two chaperone systems during proteotoxic stress.

Heat shock also redistributed IbpA toward the cell poles (**Figure 4E**), consistent with its accumulation at heat-induced protein aggregates^26,27^. IbpA was more uniformly distributed than DnaK at 37°C but became more polar during heat shock (**Figure 4F, Figure 4 – figure supplement 3A-D**). This redistribution was driven mainly by accumulation of the S_slow_ population. Thus, while DnaK remains dynamically distributed among mobility states, IbpA progressively accumulates in long-lived, low-mobility states with aggregation-prone clients. This suggested that IbpAB could buffer persistent clients and consequently preserve the availability of the DnaK pool.

To test this directly, we measured DnaK-HaloTag dynamics in Δ*ibpAB* cells. Loss of IbpAB doubled DnaK S_slow_ occupancy already at 37°C (**Figure 5A, B, Figure 5 – figure supplement 1**), indicating that IbpAB limit prolonged DnaK engagement even during steady-state growth. During heat shock, DnaK S_slow_ increased to ∼85%, and its apparent dwell time reached 1.6 ± 0.4 s after 5-7 min at 45°C (**Figure 5C**). In contrast to wild-type cells, S_slow_ occupancy showed little recovery at later time points. DnaK localization also became more polar in Δ*ibpAB* cells during heat shock (**Figure 5D, Figure 5 – figure supplement 2**), resembling the spatial distribution of IbpA (**Figure 4F**). These results support a model in which IbpAB buffer aggregation-prone clients that would otherwise cause prolonged DnaK engagement, thereby preserving DnaK availability for more rapid interactions with the proteome.

**Figure 5.**
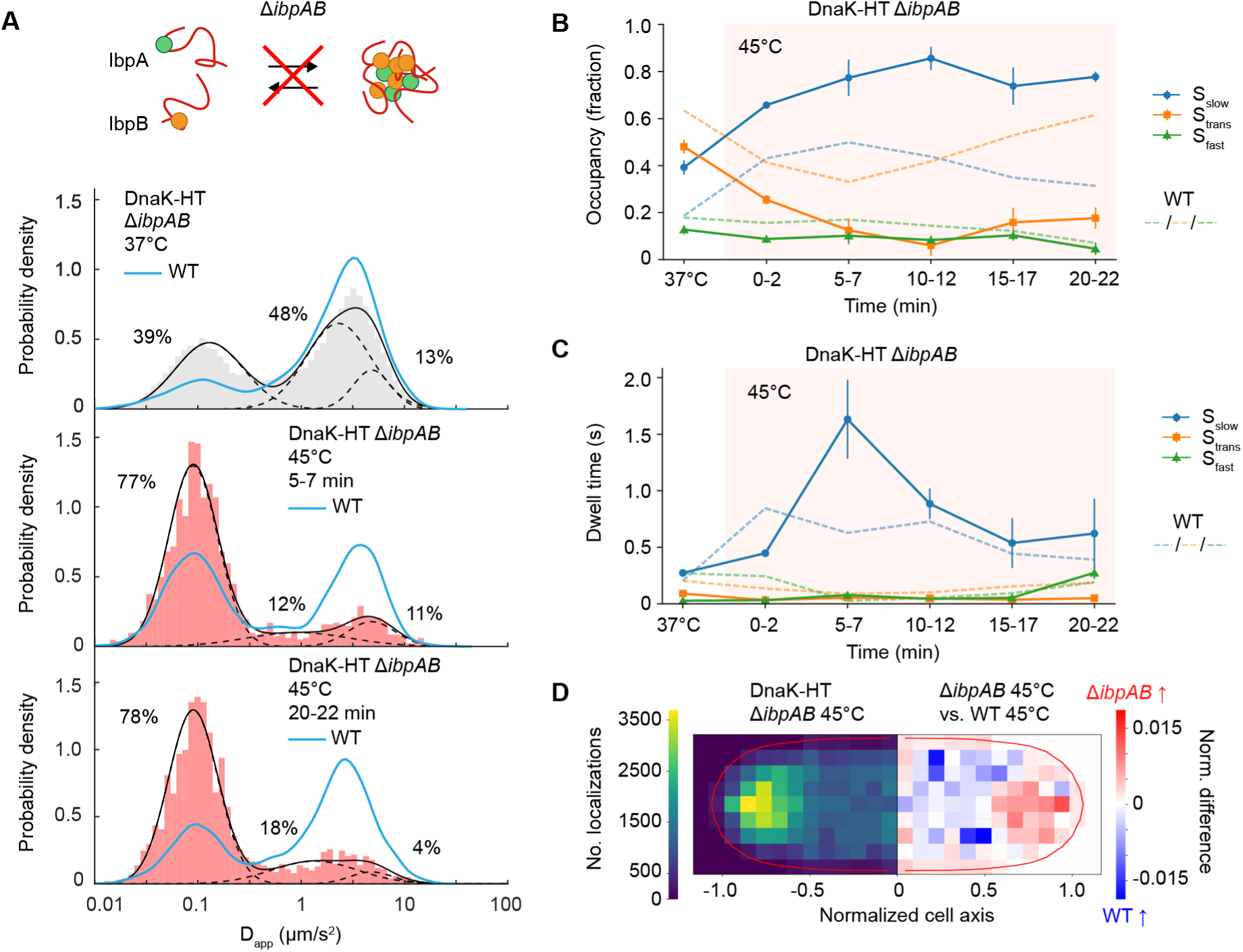
Effects of IbpAB deletion on DnaK dynamics. (**A**) Schematic representation of DnaK–HaloTag *ΔibpAB* (JML_Ec_195) and D_app_ distributions (37°C: 21946 tracks in 1139 cells; 45°C: 9423 tracks in 1467 cells); (**B**) occupancy of GMM states and (**C**) state dwell times. (**D**) Localizations of DnaK-HaloTag in *ΔibpAB* cells at 45°C (left side) and normalized differences between *ΔibpAB* and wildtype cells at 45°C. The red indicates areas of higher DnaK-HaloTag occupancy in *ΔibpAB* cells while blue pixels indicate higher DnaK occupancy in wild-type cells. Localizations are plotted on normalized cell coordinates. Data statistics are available in Supplementary Table 1-4. All data are from three biological replicates. Error bars indicate the standard error of the mean [SEM] between the replicates.

### DnaJ and HtpG modulate DnaK dynamics during sustained heat stress

To determine how other components of the chaperone network influence DnaK allocation, we analyzed DnaK dynamics in Δ*dnaJ* and Δ*htpG* cells. DnaJ promotes client transfer to DnaK and stimulates ATP hydrolysis, favoring the formation of the high-affinity client-bound state^3,34^, and has also been implicated in recruiting DnaK to protein aggregates^35^. HtpG, the bacterial Hsp90 homolog, cooperates with the DnaK system during protein folding and heat stress^36^.

Loss of DnaJ caused only a small increase in low-mobility DnaK at 37°C but led to greater accumulation of this population during prolonged heat shock, especially at later time points at 45°C (**Figure 5 – figure supplement 3A-D**). This delayed recovery is consistent with less efficient client processing or release in the absence of DnaJ. However, apparent DnaK dwell times and spatial distribution remained largely similar to wild type.

Deletion of *htpG* produced even milder effects, including a modest increase in low-mobility DnaK occupancy and a delayed increase in dwell time during heat stress (**Figure 5 – figure supplement 3A, E-G**). Thus, DnaJ and HtpG contribute to DnaK client processing primarily during sustained heat stress, but neither is required for the rapid redistribution of DnaK immediately after temperature upshift.

### Increased proteostasis demand drives heat shock-like DnaK dynamics

Cellular proteins have evolved to remain soluble within the capacity of the protein-quality-control network and have been suggested to leave little buffering capacity when protein abundance or aggregation propensity increases^37,38^. Consistent with this, overexpression of unstable or aggregation-prone proteins can induce increased production of chaperones^39,40^. We therefore asked whether the observed DnaK dynamics are specific to elevated temperature or instead reflects a more general response to increased proteostasis burden (**Figure 6A**).

**Figure 6.**
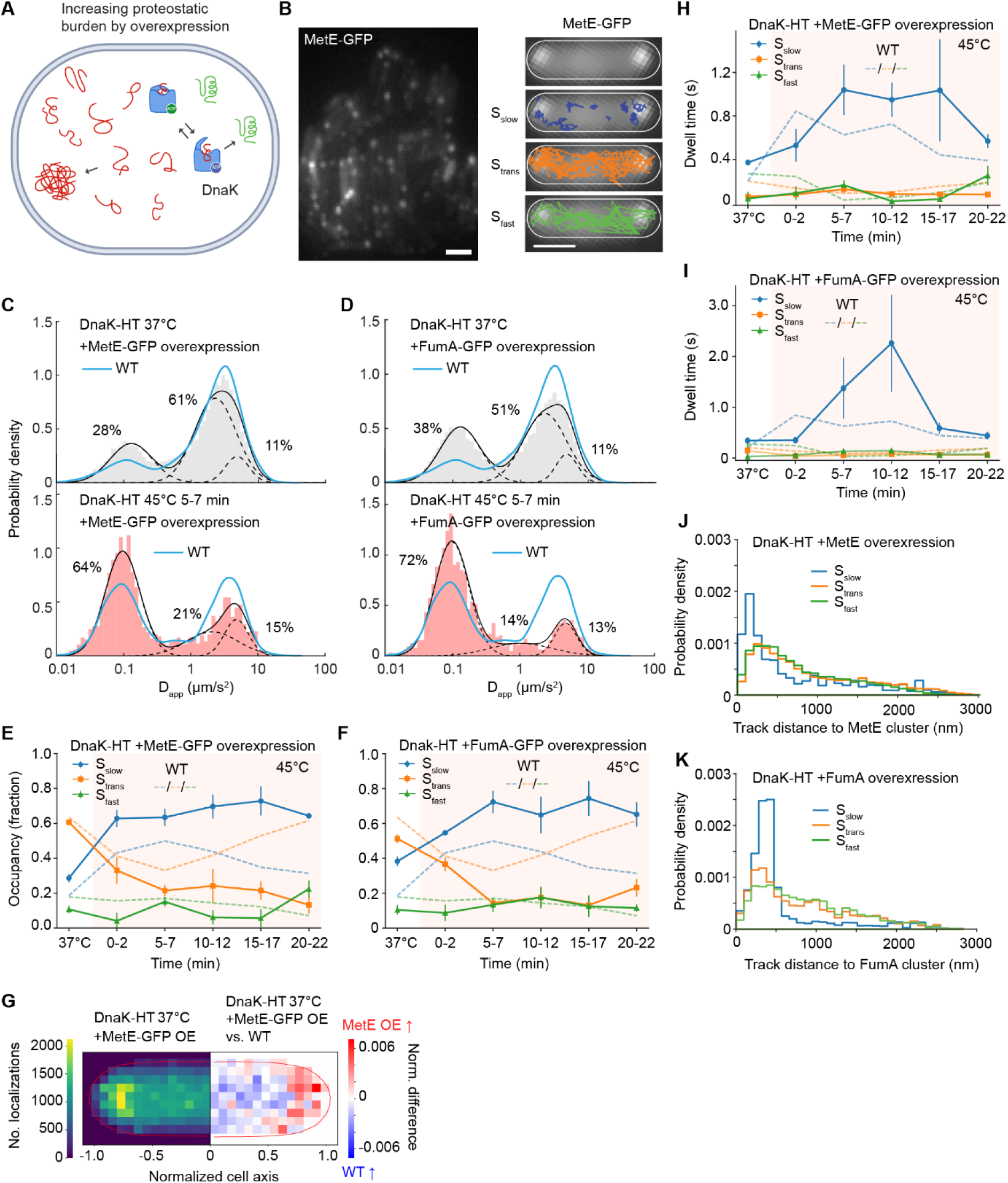
Thermolabile protein overexpression induces heat shock-like response in DnaK. (**A**) Schematic of cells overexpressing thermolabile protein substrates of DnaK. (**B**) Left: Representative fluorescence image of cells overexpressing MetE-GFP (scale bar 2 µm). Right: Representative single-molecule tracks from diffusional states of DnaK in cells overlaid with MetE fluorescence signal (scale bar 1 µm). D_app_ of DnaK-HaloTag in cells overexpressing (**C**) MetE-GFP (JML_Ec_157; 37°C: 14643 tracks in 449 cells; 45°C: 6659 tracks in 865 cells) or (**D**) FumA-GFP (JML_Ec_219; 37°C: 18276 tracks in 827 cells; 45°C: 7176 tracks in 925 cells). The D_app_ distributions were fitted with a three-state Gaussian Mixture Model (GMM). GMM-derived state occupancy for diffusional states of DnaK-HaloTag at 37°C and during 45°C heat shock in cells overexpressing (**E**) MetE-GFP or (**F**) FumA-GFP. (**G**) Localizations of DnaK-HaloTag in cells overexpressing MetE-GFP at 37°C (left side) and normalized differences between the MetE-GFP overexpression and wildtype cells at 37°C. The red indicates areas of higher DnaK-HaloTag occupancy in cells overexpressing MetE-GFP while blue pixels indicate higher DnaK occupancy in wildtype cells. Localizations are plotted on normalized cell coordinates. Hidden Markov model (HMM)-derived dwell times of the DnaK states in cells overexpressing (**H**) MetE-GFP or (**I**) FumA-GFP. Distribution of distances from a DnaK track to the closest focus of overexpressed (**J**) MetE-GFP or (**K**) FumA-GFP in each state. Data statistics are available in Supplementary Table 1-4. All data are from three biological replicates. Error bars indicate the standard error of the mean [SEM] between the replicates.

We selected two native thermolabile DnaK clients^41^, the metabolic enzymes FumA and MetE, and overexpressed them as GFP fusions from ASKA+ plasmids^42^ in the DnaK-HaloTag strain (**Figure 6B**). Induction increased MetE and FumA abundance by 1.8-fold and 2.2-fold, respectively, relative to wild-type levels with little or no detectable effect on viability after heat shock (**Figure 6 – figure supplement 1A-B**). Under these conditions, MetE-GFP and FumA-GFP formed distinct foci in 42 ± 6 % and 31 ± 4 % of cells, respectively, compared with 8 ± 1% and 10 ± 1% at endogenous expression levels (**Figure 6 – figure supplement 2**).

Overexpression of either client increased DnaK S_slow_ occupancy already at 37°C, with a stronger effect for FumA (∼2.1-fold) than MetE (∼1.5-fold) (**Figure 6C-F**). DnaK also became more enriched toward the cell poles (**Figure 6G, Figure 6 – figure supplement 3**), and S_slow_ dwell time increased (**Figure 6H, I**). As such, increased client burden alone is sufficient to shift DnaK toward more persistent low-mobility engagements at optimal growth temperature.

We next quantified whether DnaK was spatially enriched near overexpressed protein foci. For each DnaK track, we measured the distance to the nearest FumA-GFP or MetE-GFP focus as a function of DnaK mobility state (**Figure 6J, K**). S_slow_ molecules were located closest to foci, with median distances of 387 ± 44 nm for FumA-GFP and 516 ± 173 nm for MetE-GFP. In comparison, S_trans_ and S_fast_ molecules were farther from the nearest focus (FumA-GFP: 717 ± 53 nm and 778 ± 75 nm; MetE-GFP: 648 ± 106 nm and 666 ± 18 nm, respectively). Although many DnaK molecules remained outside visible foci, these results show that the low-mobility population is preferentially enriched near sites of client accumulation, indicating that DnaK is reallocated to sites of increased proteostasis demand.

Finally, we examined the combined proteotoxic effects of client overexpression and heat stress. In both FumA- and MetE-overexpressing cells, heat shock caused a strong increase in S_slow_ occupancy and depletion of S_fast_ and S_trans_ states (**Figure 6C-F, Figure 6 – figure supplement 4**). Unlike wild-type cells, however, the overexpression strains showed little recovery in S_slow_ or S_trans_ occupancy during the heat-shock time course. S_slow_ dwell times were also modestly increased, while S_trans_ and S_fast_ molecules moved closer to visible foci during heat shock (**Figure 6 – figure supplement 5A-B**).

Together, these results show that increased abundance of thermolabile DnaK clients is sufficient to produce heat shock-like changes in DnaK dynamics even at 37°C. Substrate overload therefore shifts the DnaK pool from transient proteome sampling toward longer engagements. Combining this burden with heat-induced proteome unfolding further challenges the proteostasis network and limits recovery of the pre-stress DnaK distribution. Thus, DnaK responds broadly to proteostasis demand rather than to temperature alone.

## DISCUSSION

By combining live-cell single-molecule tracking with substrate-binding mutant strains, acute heat shock, chaperone-network perturbations, and thermolabile protein overexpression, we show that Hsp70 DnaK is dynamically allocated across distinct mobility states in live *E. coli* cells. These states reveal how DnaK balances rapid proteome surveillance with longer-lived engagements during normal growth and proteotoxic stress. We note that the mobility states identified here should not be interpreted as direct nucleotide or conformational states of DnaK but instead provide an *in vivo* readout of the balance between rapidly exchanging and persistent client engagement.

A central finding of this study is that most of the DnaK pool (>80%) is not freely diffusing even during steady-state growth at 37°C (**Figure 1F**). Comparison with similarly sized, freely diffusing proteins GlcB and MetE, together with the substrate-binding-deficient DnaK^S427P^ mutant, indicates that the reduced DnaK mobility largely reflects interactions with the cellular proteome (**Figure 1D, E**). This is notable because DnaK is already highly abundant under these conditions, constituting ∼1% of total cellular protein mass and an estimated 10,000–40,000 molecules per cell^43,44^. Thus, DnaK does not appear to exist primarily as a freely diffusing reserve that is active only during stress. Instead, dynamic partitioning between mobility states allows DnaK to continuously sample the proteome and rapidly redistribute when proteostasis burden increases.

Rapid and transient interactions can represent a functional mode of target search inside a cell^45–47^. Low-affinity hydrophobic motifs are normally buried in folded proteins but become exposed during synthesis, unfolding, or misfolding^8,48^. Efficient client surveillance therefore requires a balance between mobility and engagement: DnaK must exchange rapidly enough to sample the proteome broadly, while also remaining bound long enough to support productive client folding. Contrary to DNA-binding proteins^49^, intermittent-search theory predicts no universal optimal state time or ratio, but rather a balance shaped by diffusion, client accessibility, recognition kinetics, and productive engagement probability^50^.

DnaK mobility states were also spatially organized (**Figure 1I-K**). Damaged and aggregation-prone proteins can accumulate at defined subcellular regions, particularly the cell poles and nucleoid-free regions^26–28^. DnaK appears to follow the spatial distribution of its clients, linking chaperone activity not only to client identity and abundance, but also to intracellular position. Heat shock further enhanced this spatial localization (**Figure 2D-E**). Such localization could arise from nucleoid exclusion, aggregate formation, local crowding or membrane-proximal protein damage and is consistent with previous observations of polar enrichment of protein-quality-control factors^26–28^. Proteostasis demand therefore influences both the kinetics and intracellular distribution of DnaK.

Acute heat shock revealed how the pre-existing DnaK pool responds to increased folding demand. DnaK mobility changed within seconds of the temperature upshift, before accumulation of newly synthesized heat-shock proteins (**Figure 2A**). The early response was dominated by stabilization of the low-mobility state: apparent dwell time increased approximately 4-fold within 2 min, whereas entry into this state increased more gradually and reached an approximately 2-fold maximum after 10–12 min (**Figure 3B-C**). Thus, the initial redistribution of DnaK is driven primarily by prolonged molecular engagements rather than an increased capture rate. Acute proteotoxic stress can therefore rapidly reduce DnaK availability for new clients.

The distinct dynamics of IbpA (**Figure 4**) and the effects of *ibpAB* deletion (**Figure 5**) reveal how the broader proteostasis network influences DnaK availability. During heat shock, IbpA accumulated in a long-lived, low-mobility population, consistent with association with aggregation-prone proteins^11,12,14^. In contrast, loss of IbpAB doubled low-mobility DnaK occupancy already at 37 °C and caused much stronger and more persistent DnaK engagement during heat shock (**Figure 5B**). DnaK also became more localized toward the poles in Δ*ibpAB* cells (**Figure 5D**). Together, these findings support a model in which IbpAB buffer aggregation-prone clients that would otherwise impose prolonged engagements on DnaK, thereby preserving DnaK pool availability.

Other components of the chaperone network had more selective effects. Despite its canonical role in client delivery and ATP hydrolysis in the Hsp70 cycle^3,6^, loss of DnaJ did not prevent the rapid heat-induced redistribution of DnaK, but increased low-mobility occupancy during prolonged stress (**Figure 5 – figure supplement 3A-D**). This relatively mild phenotype may reflect compensation by the alternative J-domain proteins CbpA and DjlA, which have been shown to compensate for the loss of DnaJ *in vitro* and *in vivo*^51,52^. Deletion of *htpG* produced still smaller changes (**Figure 5 – figure supplement 3A, E-G**), consistent with HtpG acting primarily during later remodeling of a subset of DnaK-associated clients rather than controlling global client recognition^11,53^. DnaJ and HtpG therefore modulate DnaK dynamics during sustained stress but are not major determinants of its initial response to temperature upshift.

Thermolabile protein overexpression provided an orthogonal way to increase proteostasis demand without changing temperature. Increased FumA or MetE abundance shifted DnaK towards the low-mobility population already at 37°C, increased apparent dwell time and altered DnaK localization (**Figure 6A-I**). Low-mobility DnaK was also enriched near visible FumA or MetE foci, linking this state to regions of elevated folding demand (**Figure 6J-K**). Combining client overexpression with heat shock produced persistent DnaK redistribution with little recovery toward the pre-stress state. Thus, the changes observed during heat shock are not simply a direct consequence of temperature: elevated client burden is itself sufficient to remodel DnaK dynamics.

Together, our results support a model in which DnaK dynamically probes the proteome to capture unfolded clients rather than functioning as a passive reserve activated mainly during stress. During balanced growth, much of the DnaK pool continuously samples the proteome, while increasing folding demand shifts molecules toward prolonged client engagement. This provides a rapid response to proteotoxic stress but simultaneously reduces the fraction of DnaK immediately available for additional clients. Auxiliary systems, such as IbpAB, help preserve DnaK capacity by buffering clients that would otherwise sequester DnaK. Effective proteostasis capacity therefore depends not only on chaperone abundance, but also on how rapidly the existing chaperone pool can be redistributed and recycled across a changing proteome. Given the conserved architecture and ATP-dependent client-binding cycle of Hsp70 chaperones, these principles are likely broadly applicable to other organisms, while still being shaped by specific co-chaperone networks^5,54^.

## MATERIALS AND METHODS

### Bacterial strains and growth conditions

All bacterial strains are listed in **Supplementary Table 5** along with their sources. All strains are derivatives of *E. coli* K12 MG1655 and resistance gene cassette inserts conferring antibiotic resistance are designated *kan*, *amp* and *cm* for kanamycin (Km^r^), ampicillin (Amp^r^) and chloramphenicol (Cm^r^), respectively. The resistance genes are flanked by FLP recognition target (*frt)* sites, where mentioned, which enable excision of the inserts when the enzyme Flp recombinase is expressed from the pCP20 vector^55^. Oligonucleotides used for polymerase chain reaction are listed in **Supplementary Table 6**.

Chromosomal fusions and gene deletions were constructed using λ-RED recombination, Gibson assembly, or P1 transduction^55–57^. HaloTag along with *kan*-cassette were inserted into the native chromosomal locus using λ-RED recombination and transferred to a recipient MG1655 strain using P1 phage transduction. Chromosomal deletions were introduced using λ-RED recombination by replacing the target gene with *kan*-cassette. All bacterial strains were confirmed by both PCR and Sanger sequencing. The single-site DnaK mutants were first cloned into a plasmid vector together with the HaloTag-kanamycin resistance cassette using Gibson assembly and then amplified and inserted into the native *dnaK* locus using λ-RED recombination.

For microscopy, cells were cultured overnight in M9 minimal medium supplemented with 0.2% glycerol (v/v), 0.1% casamino acids (w/v), and 1 µg/ml thiamine (M9glyCAAT), at 30°C or 37°C. The following day, cells were diluted 1000-fold and grown to optical density (OD_600_) ∼0.1. The HaloTag was labeled with Janelia Fluor JFX554 ligand^19^, as described previously^58^. Briefly, cells were incubated with 1.5 μM JFX554 ligand in the growth medium for 30 min with shaking on a heat block, washed four times with growth medium while maintaining temperature in a heated centrifuge (Sigma 2-16KHL), and allowed to recover for 60 min (while remaining in exponential growth phase) before a final wash prior to imaging. Finally, cells were spotted on a 1% (w/v) gellan gum pad (#J63423.30, Thermo Fisher Scientific) prepared with the M9 medium and covered with a VAHEAT coverslip. For the overexpression experiments with strains JML_Ec_219 and JML_Ec_157 harboring ASKA+ plasmids (**Supplementary Table 7**), cells were additionally induced with 0.1 mM IPTG for 2 h prior to imaging.

### Colony-forming units

To determine cell viability following heat shock, a colony-forming unit (CFU) assay was performed. Briefly, overnight cultures were diluted ∼1000-fold and grown to an early exponential phase. Each culture was heat shocked at 51°C for 15 minutes. Thereafter, pre- and post-heat shock cultures were serially diluted and cells were spotted onto LB-agar plates and grown overnight at 30°C. Strains containing ASKA+ plasmids were incubated with 0.1 mM of IPTG for 2 h before heat shock.

### Optical setup

Live-cell single-molecule-tracking was performed on a custom-built total internal reflection fluorescence (TIRF) microscope built around the Rapid Automated Modular Microscope (RAMM) System (ASI Imaging) with a motorized stage, a z-motor objective mount, and a motorized filter wheel (US-2000, MIM4, TG8, ASI Imaging). Fluorescence was measured with a 100 mW 561 nm laser with 70% transmission (iChrome MLE, Toptica). The laser was collimated and focused through a 100x oil immersion objective (CFI Plan Apochromat Lambda D, NA 1.45, Nikon) onto the sample using an angle for highly inclined and laminated optical sheet illumination (HILO) (Tokunaga et al. 2008). Fluorescence emission was filtered by a dichroic mirror and notch filter (ZT405/488/561rpc and ZET405/488/561m, Chroma). Fluorescence emission was measured using an EMCCD camera (iXon Life 897, 512x512 pixels, Andor) with an EM gain of 300 and pixel size of 0.106 µm. Transmission illumination was provided by an LED source and condenser (ASI Imaging), and it was used to collect brightfield images for cell segmentation. Data were acquired with a frame time of 5.475 ms and 20,000 frames were collected for each acquisition. GFP fluorescence in appropriate strains was measured using 100 mW 488 nm laser illumination with 3% transmission (iChrome MLE, Toptica).

Temperature control and monitoring during imaging were achieved using the VAHEAT heating stage (Stable Scopes GmbH). VAHEAT chips were plasma-etched prior to the experiment to remove background fluorescent particles (Tergeo plasma cleaner, Pie Scientific). Unless otherwise stated, cells were first recorded at 30/37°C before the temperature was upshifted to 45°C. During the heat shock, samples were recorded every 5 min up to 20 min.

### Single-particle tracking and analysis of trajectories

Fluorescent spot detection and tracking were performed using Trackpy^59^, within a custom Python analysis pipeline [01_add_imaging_data.ipynb; 02_add_mask_omnipose.ipynb; 03_1ch_tracking.ipynb]. In each frame of the time-lapse movie, fluorescent spots were identified by band-pass filtering, local peak finding, and intensity thresholding. These initial localizations were subsequently refined to obtain high-precision molecular positions. Localizations detected in consecutive frames were linked into trajectories when they occurred within 1.06 µm of one another and within the same segmented cell. If multiple localizations occurred within the same linking radius, all ambiguous localizations were excluded from further analysis. Cell boundaries were segmented from brightfield images using the deep-learning-based framework, Omnipose^60^, with the [BF_model6] model. The mobility of molecules was determined by calculating the apparent diffusion coefficient for each track from the step-wise mean-squared displacement using:

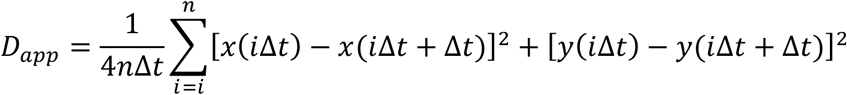

where *x(t)* and *y(t)* indicate the coordinates of the molecule at time *t*, Δ*t* is the frame interval of the camera, and *n* is the total number of the steps in the trajectory. Trajectories containing fewer than 9 displacements were omitted from the analysis, because of greater uncertainty in the calculated D_app_.

The different dynamic states of the molecules are expected to be reflected by their D_app_ values. To determine the mobility and fraction of molecules in each diffusional state, the log_10_-transformed D_app_ data of each condition were fitted to a Gaussian Mixture Model (GMM) based on the expectation-maximization algorithm^61^, as in^62^. A mixture of three Gaussian distributions contained free parameters for mean, standard deviation, and weight of each state. For IbpA-HaloTag, a three-component Gaussian mixture model with all parameters free was fitted to the experimental data. For DnaK-HaloTag data, the mean of the fast-moving fraction (S_fast_) was fixed based on the fast-moving peak of GlcB-HaloTag at each temperature (**Figure 1 – figure supplement 2A and F**). For each strain during heat shock, the mean D_app_ of the slow state (S_slow_) was fixed to a strain-specific value calculated across all heat-shock time points rather than fitted independently at each time point.

### Hidden Markov modelling

The ExTrack Hidden Markov Model (HMM) package for Python^30^ was used to analyze transition rates between the dynamic states. The algorithm was initialized with three diffusional states and the average estimates of the diffusion coefficients for each state were the same mean D_app_ values obtained from GMM fitting of the experimental data, based on the controls. For each diffusional state, the HMM algorithm inferred the average diffusion coefficient, state occupancy, and transition probabilities between each state. The average dwell was obtained by calculating the total exit rates from each state.

For some individual replicates, HMM output showed flicker or sticky dwell-time artifacts, assigning unrealistically rapid transitions out of S_fast_, determined by the software, rather than reflecting true molecular state changes. This is exacerbated by the fact that S_fast_ is a short-lived state and S_trans_ diffusion coefficient spans a broad range that overlaps with S_fast_, making dwell times sometimes difficult to resolve. To address this, the outgoing transition probabilities from the fast state (S_fast_) were constrained [extrack_constrained.py] in datasets exhibiting artifacts. The total probability of leaving S_fast_ was limited so that the model could not assign dwell times shorter than approximately three frames, which is the minimum timescale we considered reliably resolvable. A small lower bound was included for numerical stability, while the relative preference for either transition was left free to vary. This preserved flexibility in the transition model while preventing the fit from producing artifactually short S_fast_ dwell times.

### Localization analysis

2D heatmaps of single-molecule localizations were generated using the function [spatial_distribution] in [single_molecule_pipeline_python.py]. For each cell, single-molecule coordinates were projected onto normalized cell axes defined by principal component analysis of the corresponding cell-mask coordinates [sklearn.decomposition.PCA]. For each data set, the cell contour was generated by assigning the dataset’s raw mask coordinates onto the same 2D grid as the heatmap, smoothing that mask histogram with a Gaussian filter (σ = 0.5), computing an Otsu threshold (scaled by 0.5), and drawing the isocontour at this threshold. The resulting binary mask was also applied to zero out heatmap values outside the contour. The heatmaps were normalized by the total number of localizations and the difference heatmaps were computed by subtracting the normalized heatmaps from each other. Cells larger than 2.5 µm^2^ were omitted from the analysis to avoid biasing the data due to the presence of multiple nucleoids in long cells. The script also produces 1D distributions (histogram and kernel density estimate outline) along cell length. Additionally, it calculates relative densities by dividing each masked heatmap by a per-dataset localization count and plots the empirical cumulative density functions per state.

Distance from DnaK tracks to clusters of MetE-GFP or FumA-GFP was analyzed using the script [distance_to_aggregates.py]. Clusters were detected using Trackpy and in cases where cells contained several clusters, the distance to the nearest cluster was used for the subsequent analysis. Fluorescence intensity per cell was analyzed using [intensity_per_cells.py].

## Data availability

The microscopy images and data frames used in this study are available on BioStudies and the BioImage Archive (accession #XXX). Strains generated in this study are available upon request.

## Code availability

The analysis codes are available at: https://github.com/Makelalab/Published/tree/main/pajari_et_al_2026

## Acknowledgements

We thank other members of the Mäkelä group for their support, discussion, and critical reading of the manuscript. We are grateful for gifts of JFX554, a DnaKV436F plasmid, and an Omnipose model from Dr. Luke Lavis, Dr. Victor Sourjik, and Dr. Paul Wiggins, respectively. This work was supported by the European Research Council (ERC StG TEMPADAPT, 101075984) and the Research Council of Finland (349116 and 353572). A.P. was supported by the University of Helsinki Microbiology and Biotechnology doctoral programme grant. The computations and data handling were enabled by resources provided by the CSC – IT Center for Science, with the technical assistance from Susanne Merz and Simo Tuomisto (Research Software Engineer program, Aalto Scientific Computing). The funders had no role in study design, data collection and interpretation, or the decision to submit the work for publication.

## Author contributions

Conceptualization, A.P. and J.M.; Methodology, A.P.; Investigation, A.P. and J.M.; Funding acquisition, J.M.; Formal analysis A.P., Data curation A.P.; Visualization A.P. and J.M.; Project administration, J.M; Software A.P. and T.R.; Supervision, J.M. and T.R.; Writing – original draft, A.P. and J.M.; and Writing – review & editing, A.P., T.R., and J.M.

---

**Figure 1-figure supplement 1.**
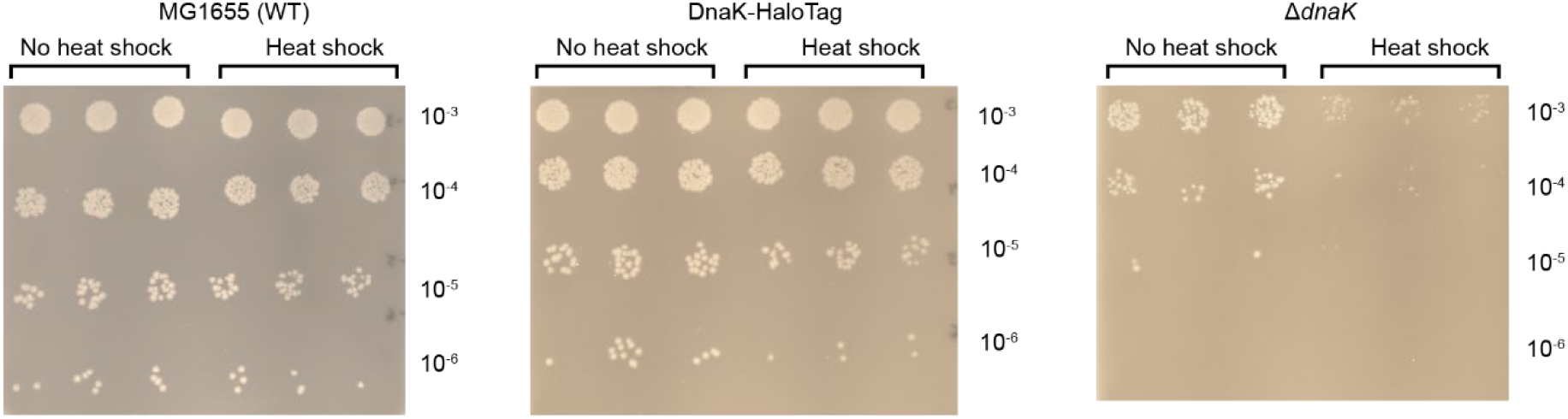
Colony forming unit assay of DnaK-HaloTag (JML_Ec_05), wild-type (MG1655) and the *ΔdnaK* strain (JML_Ec_60).

**Figure 1-figure supplement 2.**
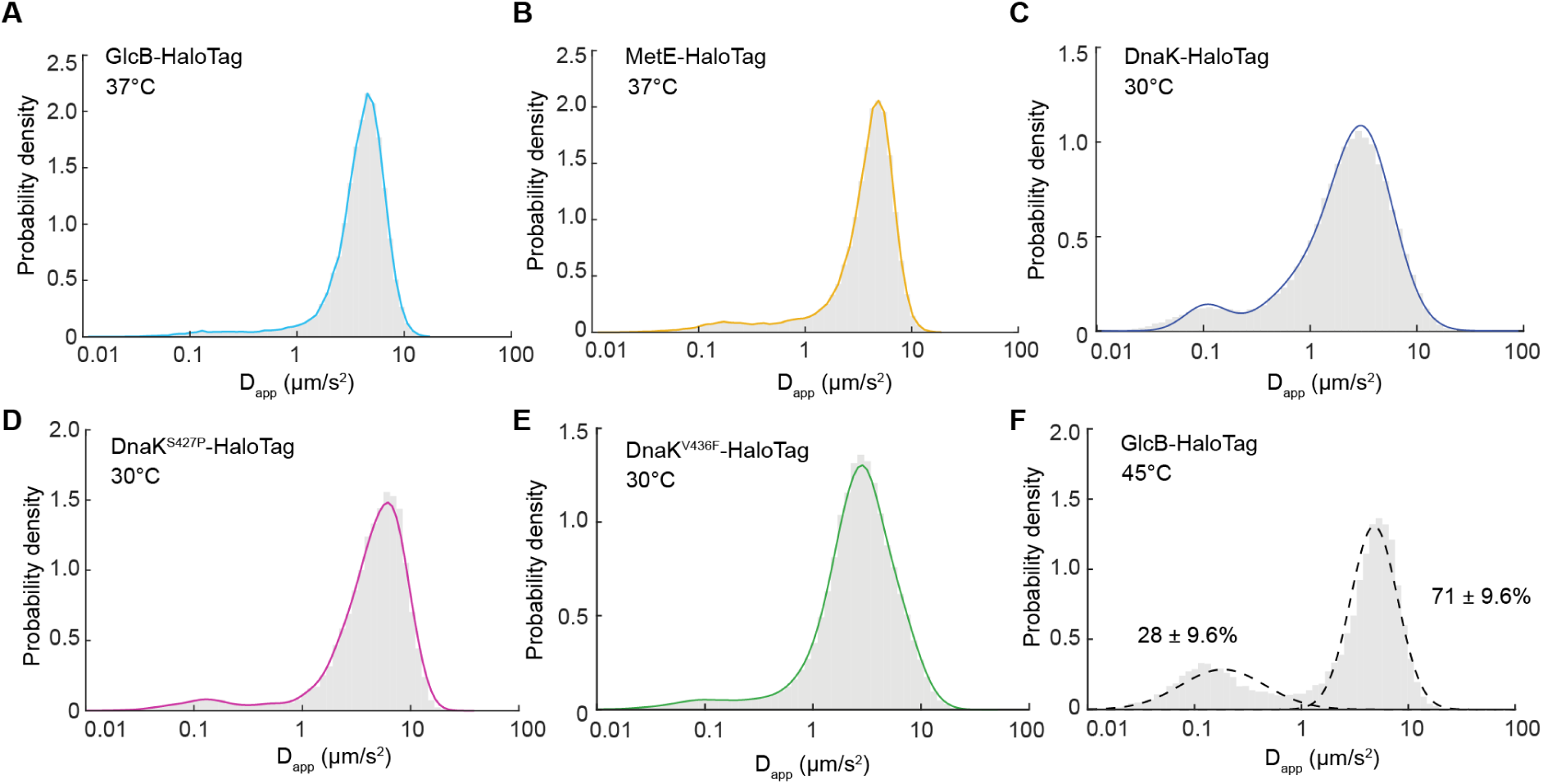
(**A**) Apparent diffusion coefficient (D_app_) distribution of JFX554-labeled GlcB-HaloTag (JML_Ec_141) at 37°C. (**B**) D_app_ distribution of JFX554-labeled MetE-HaloTag (JML_Ec_136) at 37°C. (**C**) D_app_ distribution of JFX554-labeled DnaK-HaloTag (JML_Ec_05) at 30°C (77994 tracks from 962 cells). (**D**) The D_app_ distribution of DnaK^S427P^-HaloTag at 30 °C. (**E**) The D_app_ distribution of DnaK^V436F^-HaloTag at 30°C (mean D_app_ 3.37 ± 0.10 µm^2^/s; 70019 tracks from 598 cells). (**F**) D_app_ distribution of JFX554-labeled GlcB-HaloTag at 45°C (23519 tracks from 857 cells). The mean D_app_ is 3.94 ± 0.46 µm^2^/s. The D_app_ distribution was fit with a two-state Gaussian mixture model. The mean D_app_ for the slow and fast diffusion states: 0.18 ± 0.24 µm^2^/s and 4.79 ± 1.35 µm^2^/s, respectively. Data are from three biological replicates. ± indicates the standard error of the mean [SEM] between the replicates.

**Figure 1-figure supplement 3.**
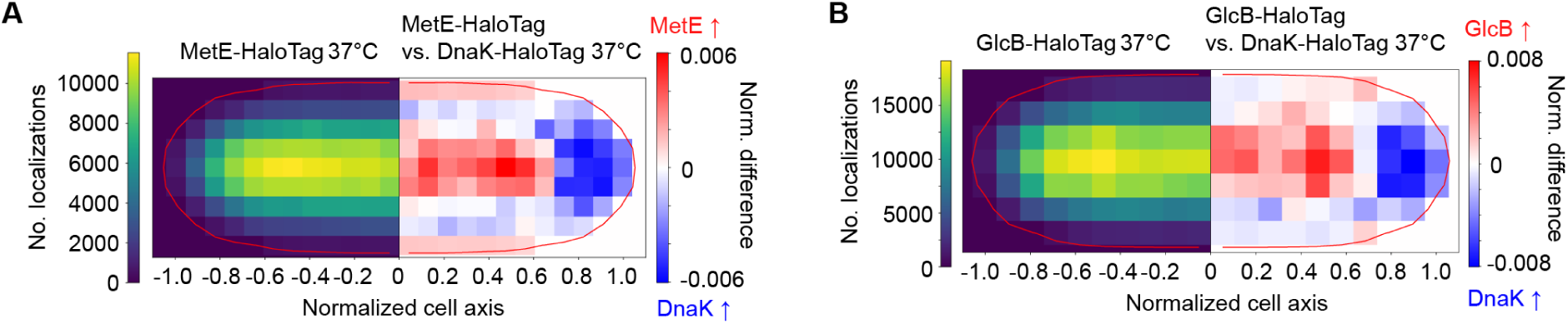
(**A**) MetE-HaloTag localizations at 37 °C (left side) and normalized difference between MetE-HaloTag and DnaK-HaloTag distributions (right side). (**B**) GlcB-HaloTag localizations at 37 °C (left side) and normalized difference between GlcB-HaloTag and DnaK-HaloTag distributions (right side). Localizations are plotted on normalized cell coordinates. Data are from three biological replicates.

**Figure 2-figure supplement 1.**
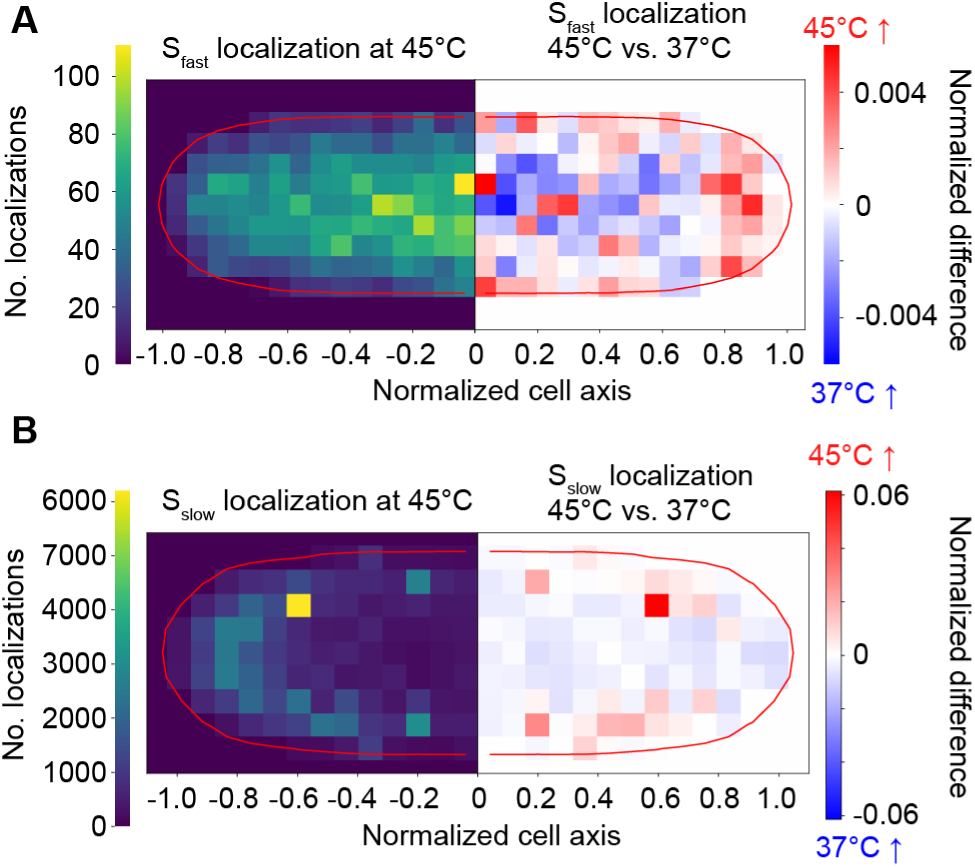
DnaK–HaloTag 45°C localization for (**A**) fast (S_fast_) and (**B**) slow (S_slow_) states. Left: distribution at 45°C; right: 45°C vs 37°C normalized difference (red = higher occupancy at 45°C, blue = higher occupancy at 37°C). Localizations are plotted on normalized cell coordinates. Data are from three biological replicates.

**Figure 3-figure supplement 1.**
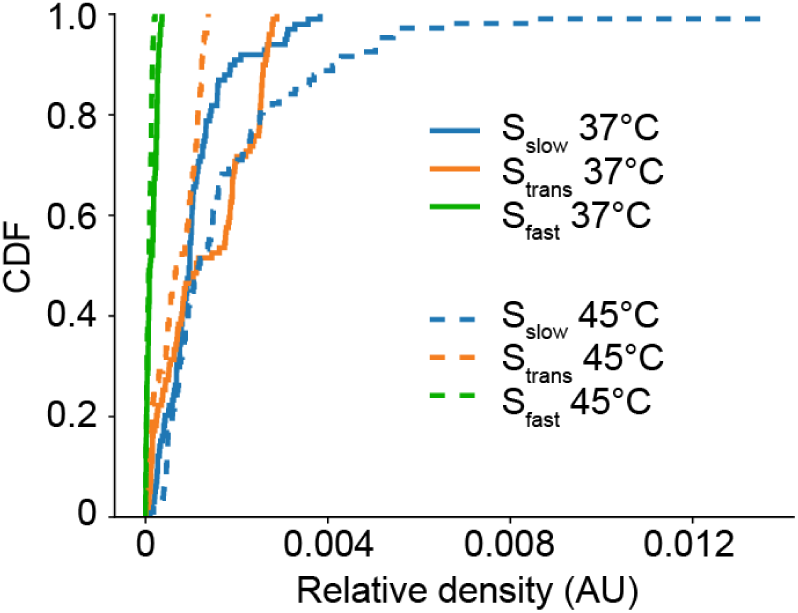
Empirical cumulative distribution functions (CDF) of the relative localization density of DnaK molecules within each state at 37°C and during 45°C heat shock. Data are from three biological replicates.

**Figure 4-figure supplement 1.**
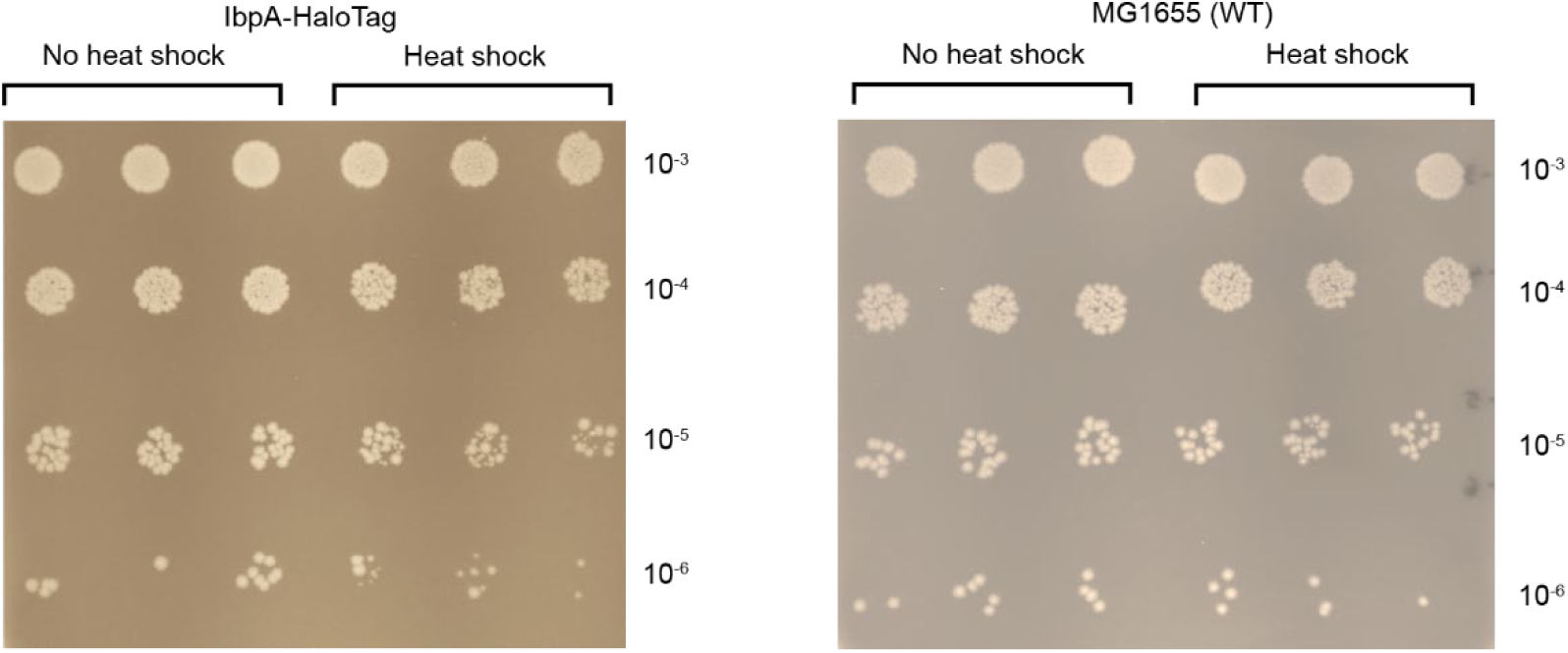
Colony forming unit assay of IbpA-HaloTag (JML_Ec_14) and wild-type (MG1655).

**Figure 4-figure supplement 2.**
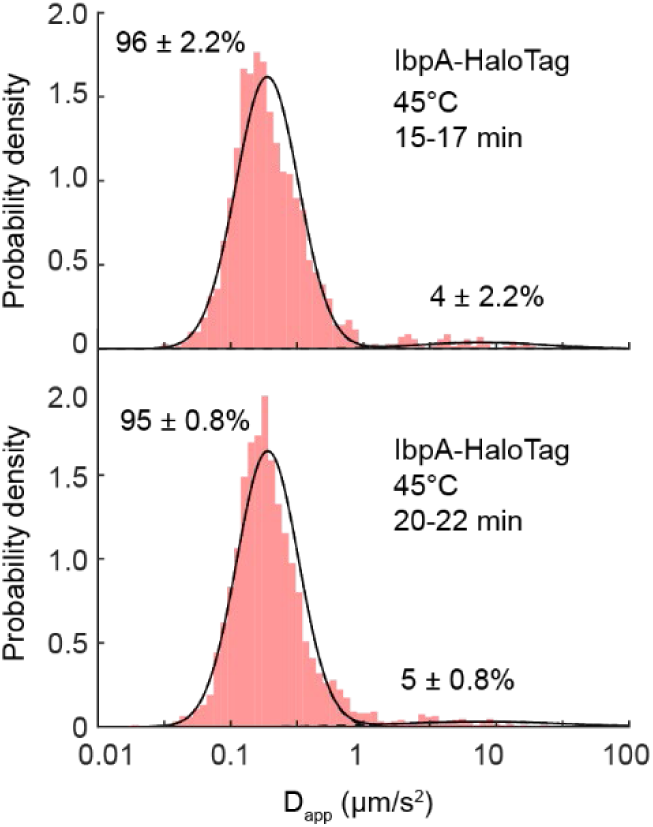
D_app_ distributions of JFX554-labeled IbpA-HaloTag at 45°C. The distributions were fit with a two-state Gaussian mixture model. The mean D_app_ for the slow diffusion state 15-17 min: 0.19 ± 0.03 µm^2^/s (1419 tracks from 166 cells); 20-22 min: 0.19 ± 0.02 µm^2^/s (2380 tracks from 183 cells). Data are from three biological replicates. ± indicates the standard error of the mean [SEM] between the replicates.

**Figure 4-figure supplement 3.**
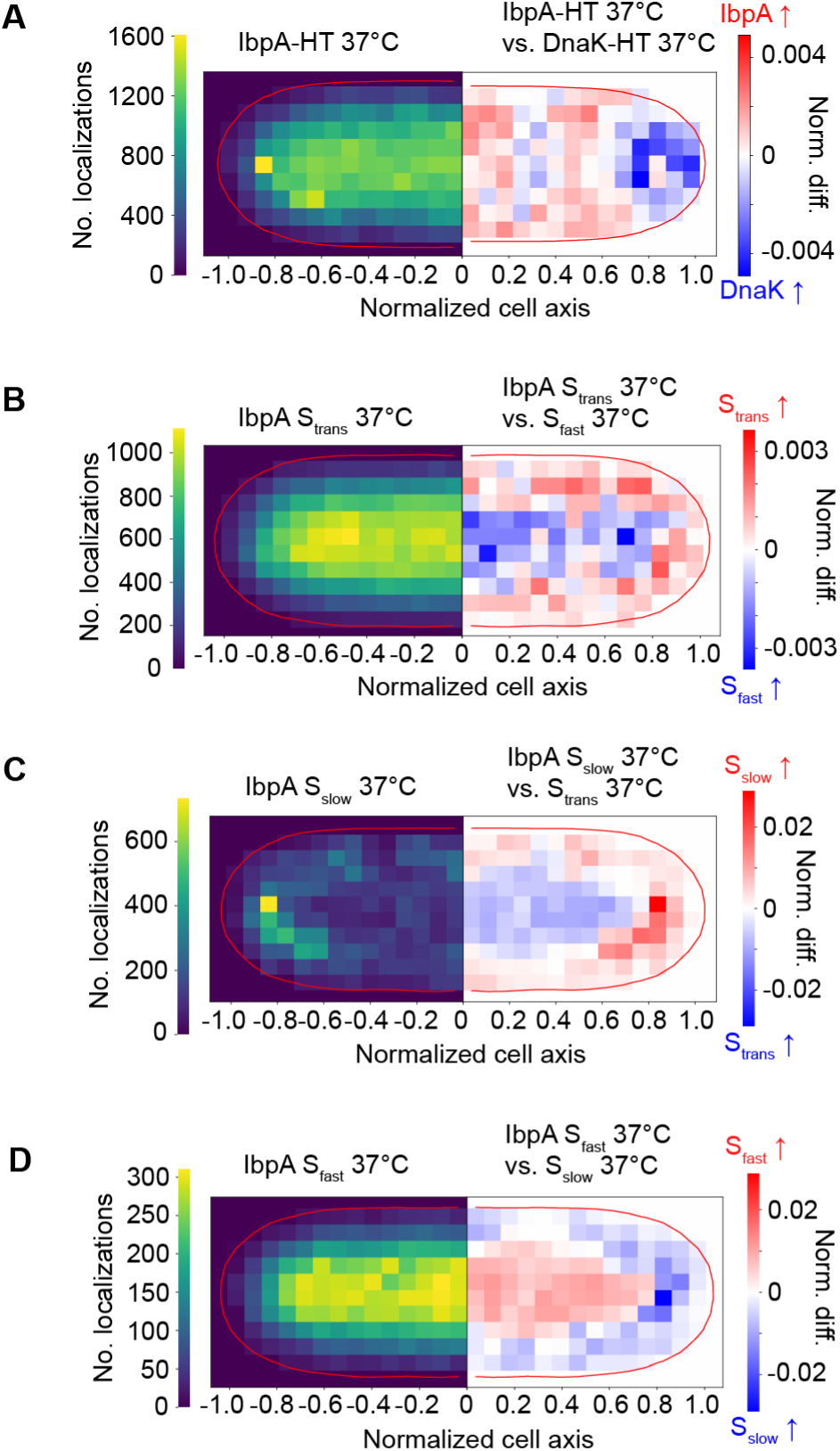
(**A**) IbpA-HaloTag localizations at 37 °C (left side) and normalized difference between IbpA localizations and DnaK-HaloTag localizations at 37 °C (right side). (**B-D**) IbpA localizations at 37 °C of diffusional states S_slow_, S_trans_ and S_fast_ (left side) and normalized differences between the states (right side). Localizations are plotted on normalized cell coordinates. Data are from three biological replicates.

**Figure 5-figure supplement 1.**
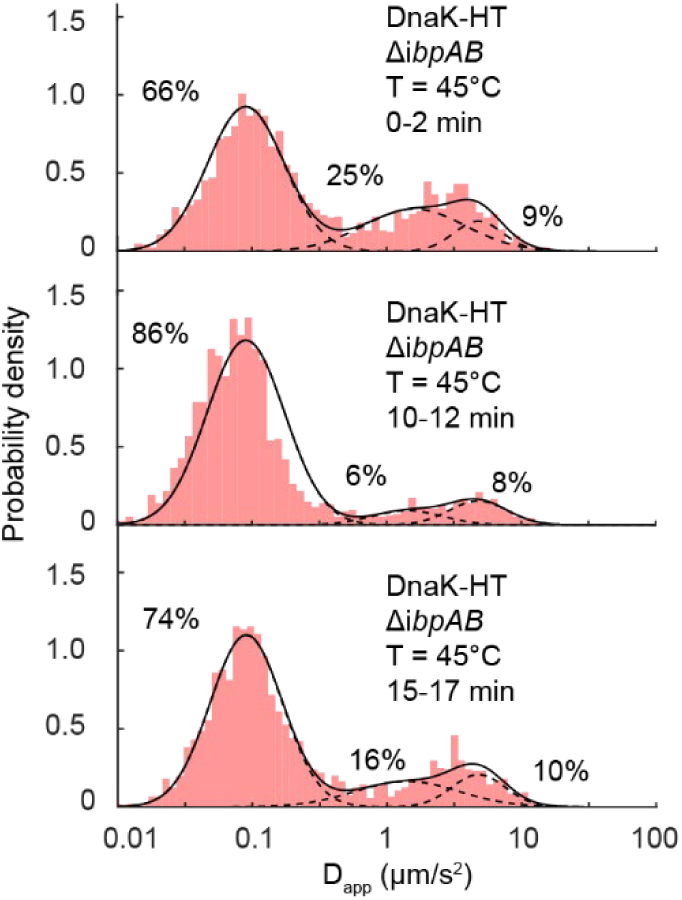
D_app_ distributions of JFX554-labeled DnaK-HaloTag in Δ*ibpAB* strain (JML_Ec_195). Data are from three biological replicates.

**Figure 5-figure supplement 2.**
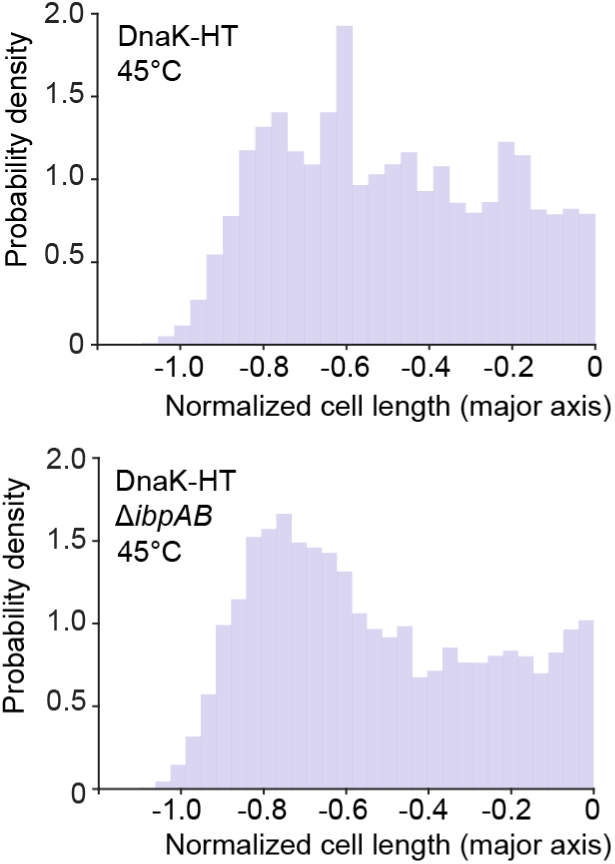
Localization distribution along normalized cell length of wild-type DnaK-HaloTag at 45°C (top) and the Δ*ibpAB* strain (bottom). Data are from three biological replicates.

**Figure 5-figure supplement 3.**
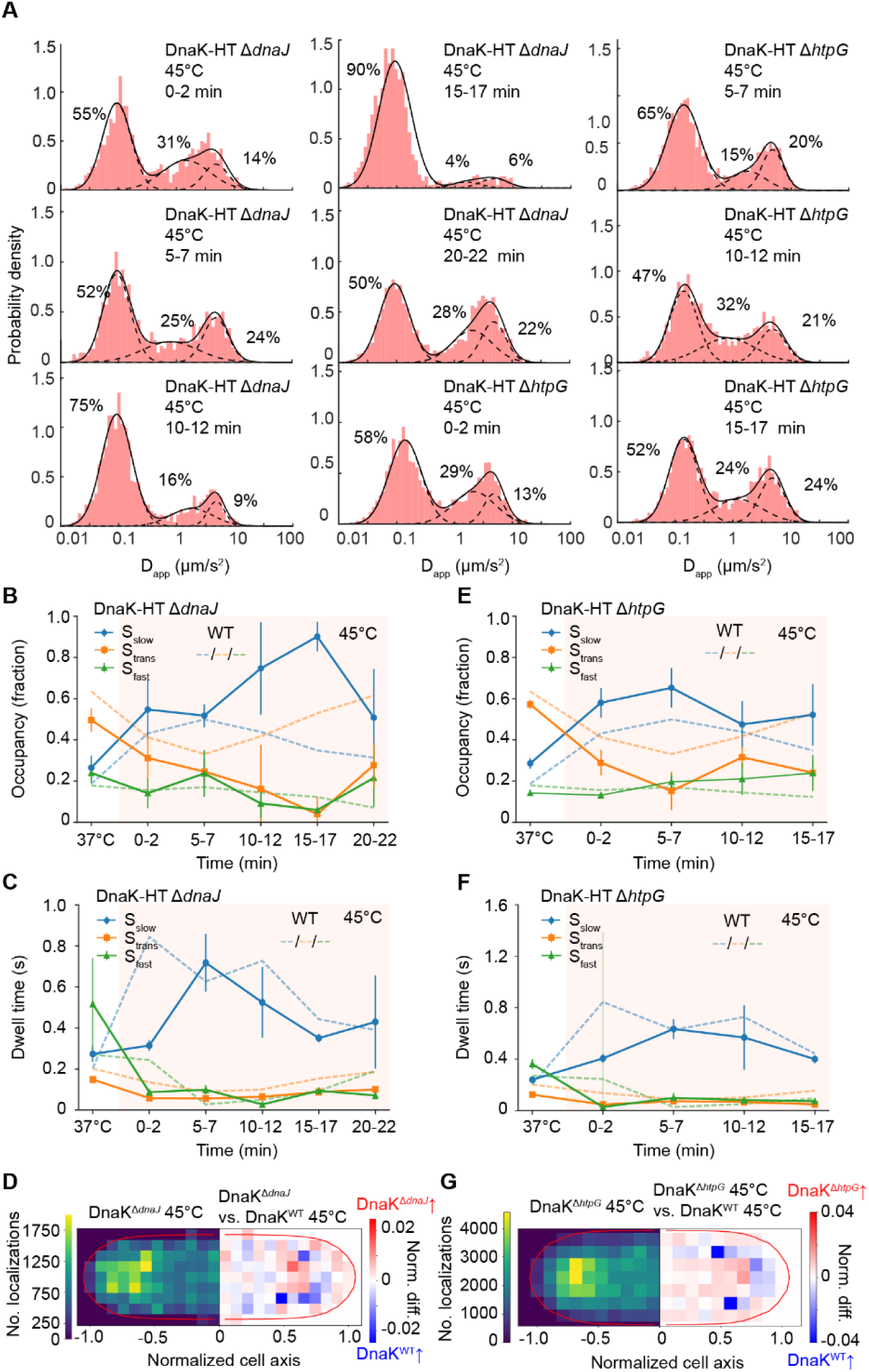
(**A**) D_app_ distributions of JFX554-labeled DnaK-HaloTag in co-chaperone deletion strains at 45°C. The distributions were fit with a three-state Gaussian mixture model (*ΔdnaJ,* JML_Ec_202: 37°C: 12317 tracks in 724 cells; 45°C: 7747 tracks and 961 cells), *ΔhtpG,* JML_Ec_147: 37°C; 23410 tracks in 849 cells; 45°C: 10415 tracks in 1038 cells). *ΔdnaJ:* (**B**) occupancy of GMM states and (**C**) dwell times. (**D**) Localizations of DnaK-HaloTag in cells with the *ΔdnaJ* at 45°C (left) and normalized differences between *ΔdnaJ* and wildtype cells at 45°C (right). The red indicates areas of higher DnaK-HaloTag occupancy in *ΔdnaJ* cells while blue pixels indicate higher DnaK occupancy in wildtype cells. *ΔhtpG:* (**E**) occupancy of GMM states and (**F**) dwell times. (**G**) Localizations of DnaK-HaloTag in cells with the *ΔhtpG* at 45°C (left) and normalized differences between *ΔhtpG* and wildtype cells at 45°C (right). Localizations are plotted on normalized cell coordinates. Data statistics are available in Supplementary Table 1-4. Data are from three biological replicates. Error bars indicate the standard error of the mean [SEM] between the replicates.

**Figure 6-figure supplement 1.**
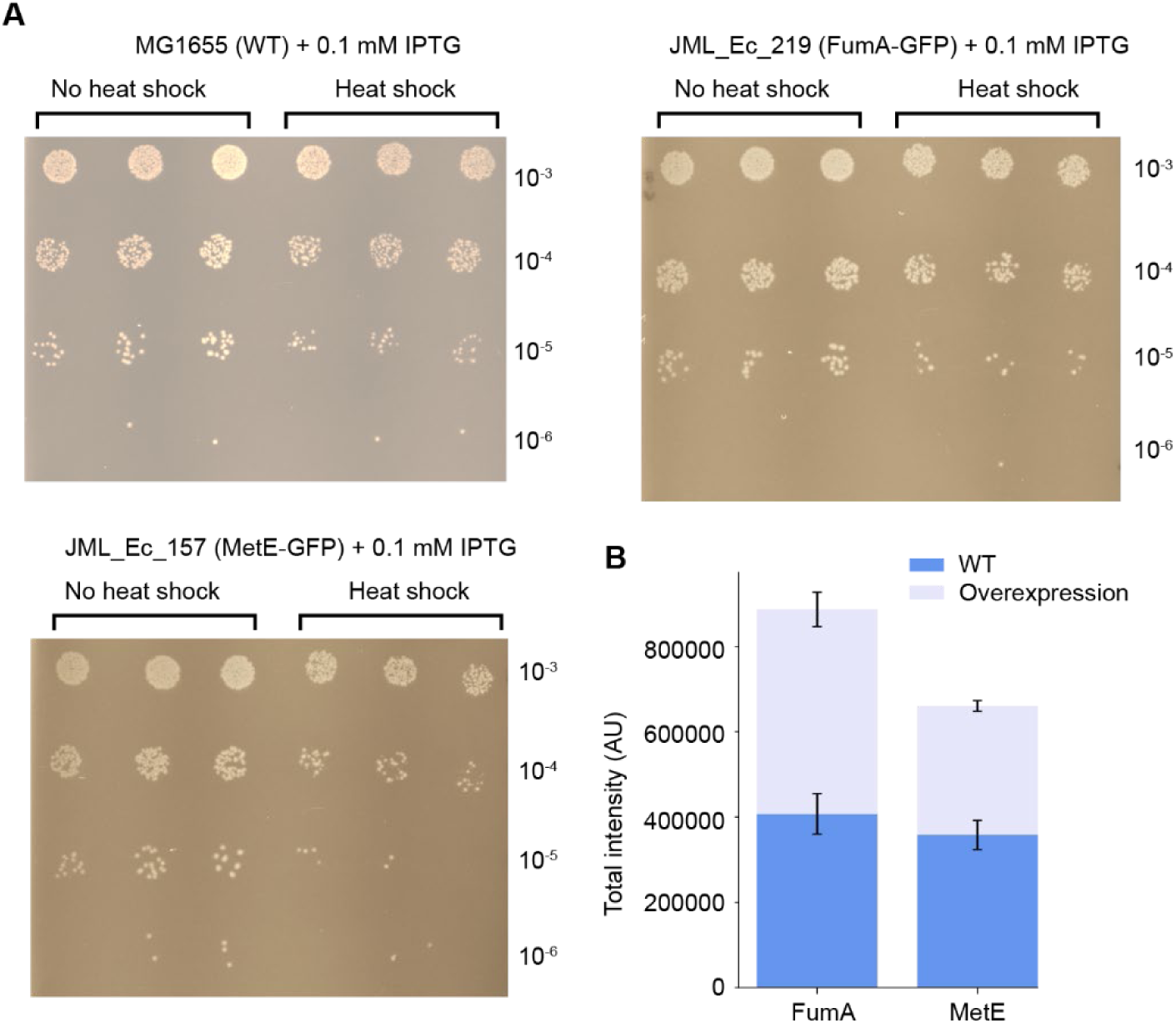
(**A**) Colony forming unit assay of strains harbouring an ASKA+ overexpression plasmid for either FumA-GFP (JML_Ec_219) or MetE-GFP (JML_Ec_157) and wild-type background (MG1655). All three cultures were induced with 0.1 mM IPTG for 2 h prior to measurements. (**B**) Fluorescence intensity in WT cells (blue) expressing FumA-GFP (JML_Ec_250, 3015 cells) or MetE-GFP (JML_Ec_249, 2870 cells) and in cells transformed with an ASKA+ overexpression plasmid (gray) for either MetE-GFP (JML_Ec_157) or FumA-GFP (JML_Ec_219). Strains containing an overexpression plasmid were induced using 0.1 mM IPTG for 2 h prior to imaging. Error bars denote the standard error of the mean [SEM] between the three biological replicates.

**Figure 6-figure supplement 2.**
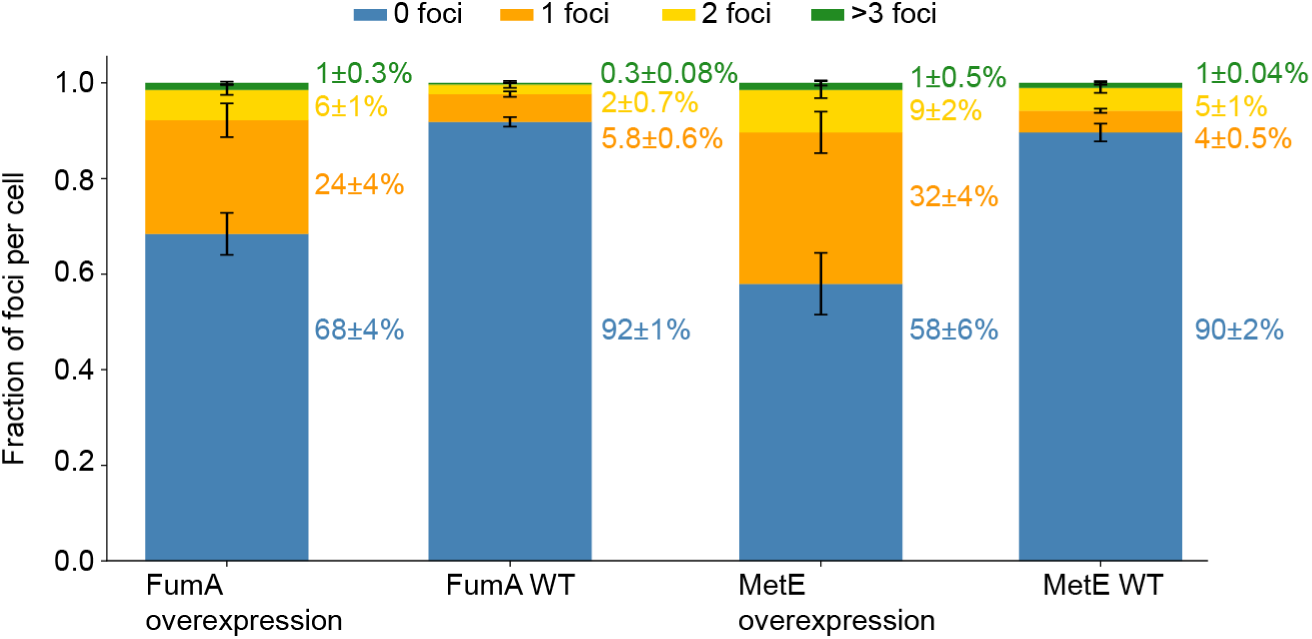
Fraction of cells with 0, 1, 2, or ≥3 FumA-GFP or MetE-GFP foci. Error bars and ± denote the standard error of the mean [SEM] between the three biological replicates.

**Figure 6-figure supplement 3.**
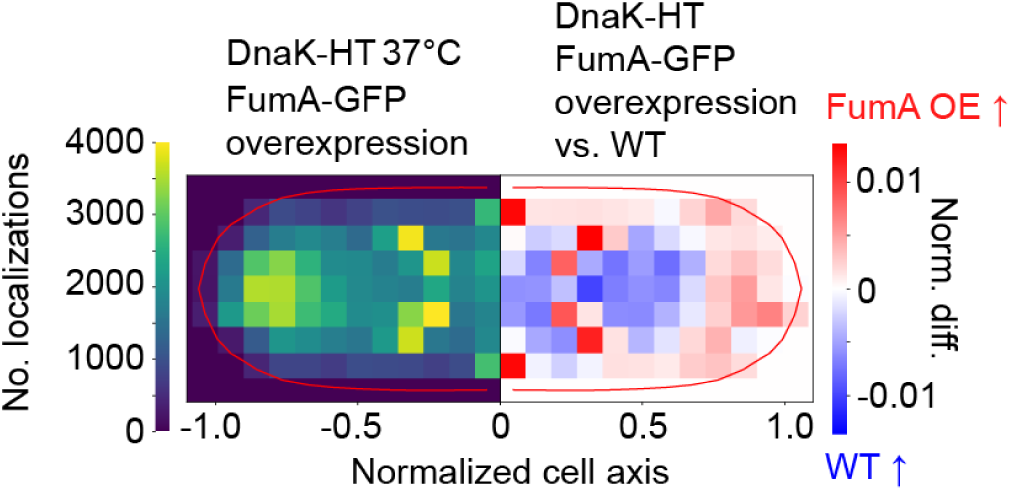
DnaK-HaloTag localization in FumA-GFP overexpression strain (JML_Ec_219) at 37 °C (left) and relative difference between FumA-GFP overexpression strain and wild-type DnaK-HaloTag distributions (right). Localizations are plotted on normalized cell coordinates. Data are from three biological replicates.

**Figure 6-figure supplement 4.**
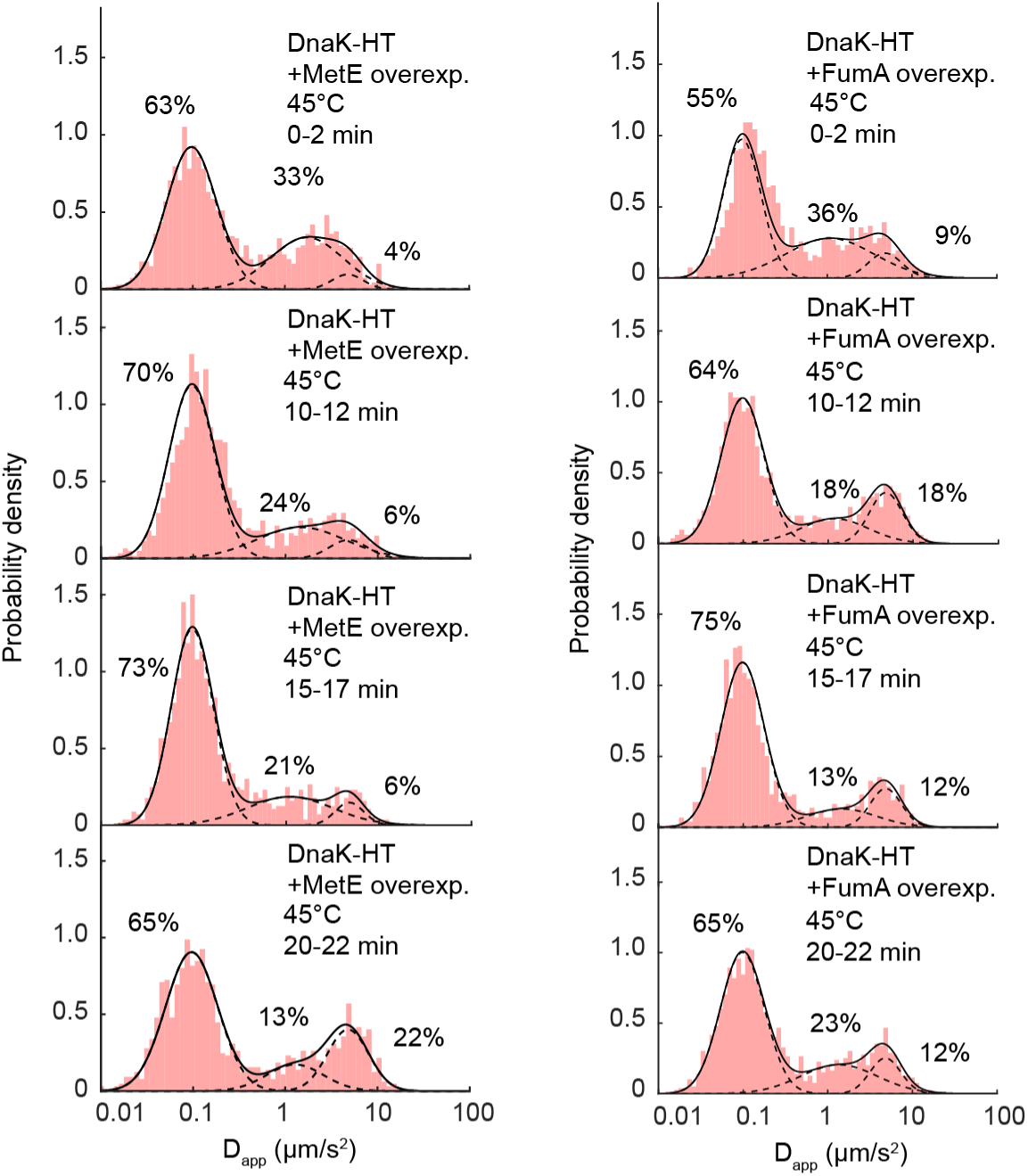
D_app_ distributions of DnaK-HaloTag in strains overexpressing MetE-GFP or FumA-GFP fitted with a three-state Gaussian Mixture Model (GMM). Data are from three biological replicates.

**Figure 6-figure supplement 5.**
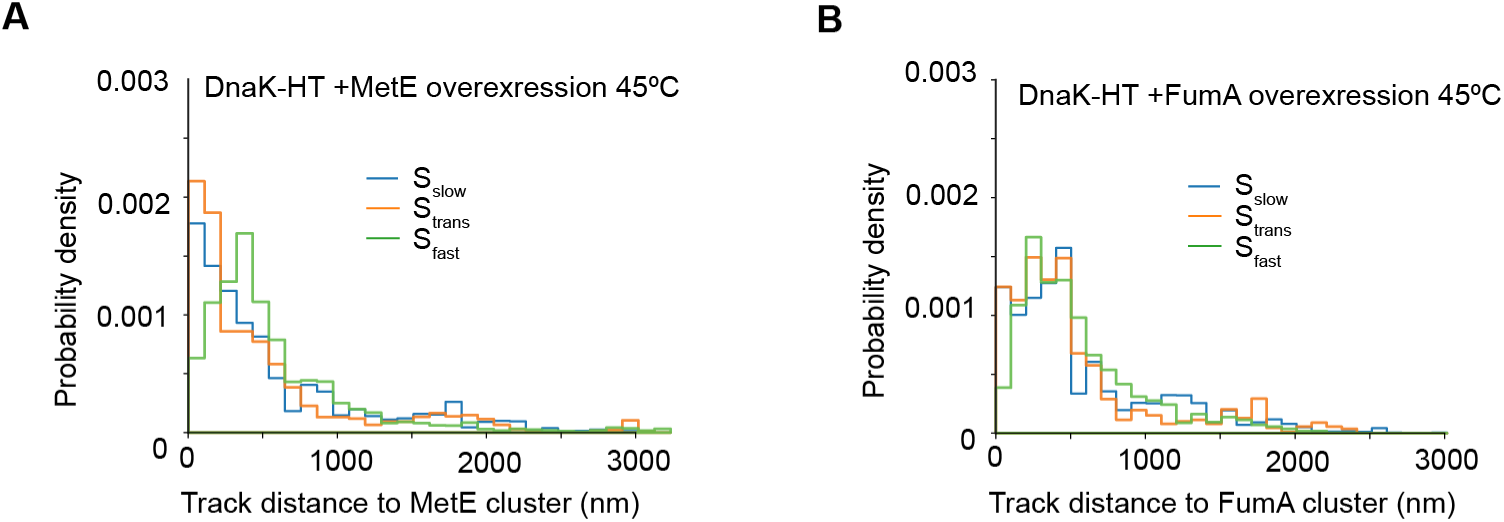
Distribution of distance from a DnaK track in each state to the closest (**A**) MetE-GFP or (**B**) FumA-GFP focus at 45°C. Data are from three biological replicates.

**Supplementary Table 1.** State occupancy values for strains at 45°C heat shock.

| DnaK-HaloTag (JML_Ec_05) state occupancy values at 45°C heat shock |  |  |
| --- | --- | --- |
| Timepoint at 45°C | Occupancy (%) ±SEM | State |
| 0-2 min | 43 ± 4 | S<sub>slow</sub> |
| 0-2 min | 41 ± 6 | S<sub>trans</sub> |
| 0-2 min | 16 ± 2 | S<sub>fast</sub> |
| 5-7 min | 50 ± 8 | S<sub>slow</sub> |
| 5-7 min | 33 ± 9 | S<sub>trans</sub> |
| 5-7 min | 17 ± 1 | S<sub>fast</sub> |
| 10-12 min | 44 ± 0.2 | S<sub>slow</sub> |
| 10-12 min | 42 ± 0.1 | S<sub>trans</sub> |
| 10-12 min | 14 ± 2 | S<sub>fast</sub> |
| 15-17 min | 35 ± 6 | S<sub>slow</sub> |
| 15-17 min | 53 ± 9 | S<sub>trans</sub> |
| 15-17 min | 12 ± 10 | S<sub>fast</sub> |
| 20-22 min | 31 ± 10 | S<sub>slow</sub> |
| 20-22 min | 62 ± 9 | S<sub>trans</sub> |
| 20-22 min | 7 ± 2 | S<sub>fast</sub> |
| IbpA-HaloTag (JML_Ec_14) state occupancy values at 45°C heat shock. |  |  |
| Timepoint at 45°C | Occupancy (%) ±SEM | State |
| 0-2 min | 34 ± 6 | S<sub>slow</sub> |
| 0-2 min | 60 ± 6 | S<sub>trans</sub> |
| 0-2 min | 6 ± 1 | S<sub>fast</sub> |
| 5-7 min | 73 ± 4 | S<sub>slow</sub> |
| 5-7 min | 18 ± 3 | S<sub>trans</sub> |
| 5-7 min | 9 ± 1 | S<sub>fast</sub> |
| 10-12 min | 76 ± 5 | S<sub>slow</sub> |
| 10-12 min | 17 ± 4 | S<sub>trans</sub> |
| 10-12 min | 7 ± 1.8 | S<sub>fast</sub> |
| State occupancy values at 45°C heat shock for Δ*ibpAB* strain (JML_Ec_195). | | |
| Timepoint at 45°C | Occupancy (%) ±SEM | State |
| 0-2 min | 66 ± 1 | S<sub>slow</sub> |
| 0-2 min | 25 ± 2 | S<sub>trans</sub> |
| 0-2 min | 9 ± 2 | S<sub>fast</sub> |
| 5-7 min | 77 ± 8 | S<sub>slow</sub> |
| 5-7 min | 12 ± 4 | S<sub>trans</sub> |
| 5-7 min | 10 ± 4 | S<sub>fast</sub> |
| 10-12 min | 86 ± 5 | S<sub>slow</sub> |
| 10-12 min | 6 ± 4 | S<sub>trans</sub> |
| 10-12 min | 8 ± 1 | S<sub>fast</sub> |
| 15-17 min | 74 ± 8 | S<sub>slow</sub> |
| 15-17 min | 16 ± 6 | S<sub>trans</sub> |
| 15-17 min | 10 ± 2 | S<sub>fast</sub> |
| 20-22 min | 78 ± 2 | S<sub>slow</sub> |
| 20-22 min | 18 ± 4 | S<sub>trans</sub> |
| 20-22 min | 5 ± 3 | S<sub>fast</sub> |
| State occupancy values at 45°C heat shock for $\Delta dnaJ$ strain (JML_Ec_202). | | |
| Timepoint at 45°C | Occupancy (%) ±SEM | State |
| 0-2 min | 55 ± 15 | S<sub>slow</sub> |
| 0-2 min | 31 ± 10 | S<sub>trans</sub> |
| 0-2 min | 14 ± 7 | S<sub>fast</sub> |
| 5-7 min | 52 ± 6 | S<sub>slow</sub> |
| 5-7 min | 25 ± 6 | S<sub>trans</sub> |
| 5-7 min | 24 ± 11 | S<sub>fast</sub> |
| 10-12 min | 75 ± 22 | S<sub>slow</sub> |
| 10-12 min | 16 ± 21 | S<sub>trans</sub> |
| 10-12 min | 9 ± 7 | S<sub>fast</sub> |
| 15-17 min | 90 ± 7 | S<sub>slow</sub> |
| 15-17 min | 4 ± 8 | S<sub>trans</sub> |
| 15-17 min | 6 ± 1 | S<sub>fast</sub> |
| 20-22 min | 51 ± 24 | S<sub>slow</sub> |
| 20-22 min | 28 ± 11 | S<sub>trans</sub> |
| 20-22 min | 22 ± 14 | S<sub>fast</sub> |
| State occupancy values at 45°C heat shock for $\Delta htpG$ strain (JML_Ec_147). | | |
| Timepoint at 45°C | Occupancy (%) ±SEM | State |
| 0-2 min | 58 ± 7 | S<sub>slow</sub> |
| 0-2 min | 29 ± 6 | S<sub>trans</sub> |
| 0-2 min | 13 ± 2 | S<sub>fast</sub> |
| 5-7 min | 65 ± 10 | S<sub>slow</sub> |
| 5-7 min | 15 ± 9 | S<sub>trans</sub> |
| 5-7 min | 20 ± 3 | S<sub>fast</sub> |
| 10-12 min | 47 ± 11 | S<sub>slow</sub> |
| 10-12 min | 32 ± 5 | S<sub>trans</sub> |
| 10-12 min | 21 ± 8 | S<sub>fast</sub> |
| 15-17 min | 52 ± 15 | S<sub>slow</sub> |
| 15-17 min | 24 ± 6 | S<sub>trans</sub> |
| 15-17 min | 24 ± 9 | S<sub>fast</sub> |
| MetE-GFP overexpression strain (JML_Ec_157) state occupancy values at 45°C heat shock. |  |  |
| Timepoint at 45°C | Occupancy (%) ±SEM | State |
| 0-2 min | 63 ± 5 | S<sub>slow</sub> |
| 0-2 min | 33 ± 8 | S<sub>trans</sub> |
| 0-2 min | 4 ± 4 | S<sub>fast</sub> |
| 5-7 min | 64 ± 5 | S<sub>slow</sub> |
| 5-7 min | 21 ± 3 | S<sub>trans</sub> |
| 5-7 min | 15 ± 3 | S<sub>fast</sub> |
| 10-12 min | 70 ± 7 | S<sub>slow</sub> |
| 10-12 min | 24 ± 10 | S<sub>trans</sub> |
| 10-12 min | 6 ± 4 | S<sub>fast</sub> |
| 15-17 min | 73 ± 8 | S<sub>slow</sub> |
| 15-17 min | 21 ± 5 | S<sub>trans</sub> |
| 15-17 min | 6 ± 5 | S<sub>fast</sub> |
| 20-22 min | 65 ± 3 | S<sub>slow</sub> |
| 20-22 min | 13 ± 5 | S<sub>trans</sub> |
| 20-22 min | 22 ± 5 | S<sub>fast</sub> |
| FumA-GFP overexpression strain (JML_Ec_219) state occupancy values at 45°C heat shock. |  |  |
| Timepoint at 45°C | Occupancy (%) ±SEM | State |
| 0-2 min | 55 ± 1 | S<sub>slow</sub> |
| 0-2 min | 36 ± 4 | S<sub>trans</sub> |
| 0-2 min | 9 ± 5 | S<sub>fast</sub> |
| 5-7 min | 72 ± 7 | S<sub>slow</sub> |
| 5-7 min | 14 ± 3 | S<sub>trans</sub> |
| 5-7 min | 13 ± 4 | S<sub>fast</sub> |
| 10-12 min | 64 ± 10 | S<sub>slow</sub> |
| 10-12 min | 18 ± 5 | S<sub>trans</sub> |
| 10-12 min | 18 ± 6 | S<sub>fast</sub> |
| 15-17 min | 75 ± 10 | S<sub>slow</sub> |
| 15-17 min | 13 ± 4 | S<sub>trans</sub> |
| 15-17 min | 12 ± 6 | S<sub>fast</sub> |
| 20-22 min | 65 ± 7 | S<sub>slow</sub> |
| 20-22 min | 23 ± 5 | S<sub>trans</sub> |
| 20-22 min | 12 ± 2 | S<sub>fast</sub> |

**Supplementary Table 2.** Number of single-molecule tracks and cells during heat shock.

| Number of single-molecule tracks and cells analyzed for DnaK-HaloTag (JML_Ec_05) strains at heat shock. |  |  |  |
| --- | --- | --- | --- |
| Time point | No. tracks | No. cells | Strain |
| 0-2 min | 2923 | 176 | JML_Ec_05 |
| 5-7 min | 2991 | 179 | JML_Ec_05 |
| 10-12 min | 2980 | 162 | JML_Ec_05 |
| 15-17 min | 4392 | 167 | JML_Ec_05 |
| 20-22 min | 5676 | 185 | JML_Ec_05 |

| Number of single-molecule tracks and cells analyzed for IbpA-HaloTag at heat shock. |  |  |  |
| --- | --- | --- | --- |
| Time point | No. tracks | No. cells | Strain |
| 0-2 min | 4539 | 294 | JML_Ec_14 |
| 5-7 min | 5978 | 305 | JML_Ec_14 |
| 10-12 min | 6152 | 281 | JML_Ec_14 |
| Number of single-molecule tracks and cells analyzed for co-chaperone deletion strains at heat shock. |  |  |  |
| Time point | No. tracks | No. cells | Strain |
| 0-2 min | 2198 | 322 | JML_Ec_195 |
| 5-7 min | 1411 | 248 | JML_Ec_195 |
| 10-12 min | 2071 | 284 | JML_Ec_195 |
| 15-17 min | 2004 | 318 | JML_Ec_195 |
| 20-22 min | 1739 | 295 | JML_Ec_195 |
| 0-2 min | 1694 | 211 | JML_Ec_202 |
| 5-7 min | 1314 | 199 | JML_Ec_202 |
| 10-12 min | 1388 | 177 | JML_Ec_202 |
| 15-17 min | 1151 | 203 | JML_Ec_202 |
| 20-22 min | 2200 | 171 | JML_Ec_202 |
| 0-2 min | 3207 | 273 | JML_Ec_147 |
| 5-7 min | 1783 | 248 | JML_Ec_147 |
| 10-12 min | 2346 | 273 | JML_Ec_147 |
| 15-17 min | 3079 | 244 | JML_Ec_147 |
| Number of single-molecule tracks and cells analyzed for thermolabile DnaK substrate overexpression strains JML_Ec_157 and JML_Ec_219. |  |  |  |
| Time point | No. tracks | No. cells | Strain |
| 0-2 min | 1293 | 171 | JML_Ec_157 |
| 5-7 min | 1835 | 195 | JML_Ec_157 |
| 10-12 min | 1188 | 159 | JML_Ec_157 |
| 15-17 min | 1158 | 149 | JML_Ec_157 |
| 20-22 min | 1185 | 191 | JML_Ec_157 |
| 0-2 min | 1492 | 206 | JML_Ec_219 |
| 5-7 min | 1225 | 163 | JML_Ec_219 |
| 10-12 min | 1564 | 173 | JML_Ec_219 |
| 15-17 min | 1387 | 186 | JML_Ec_219 |
| 20-22 min | 1508 | 197 | JML_Ec_219 |

**Supplementary Table 3.**
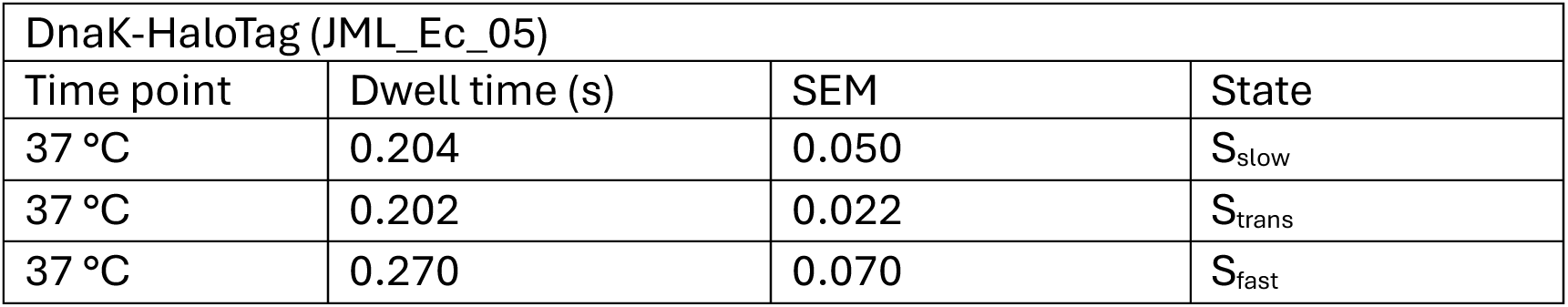
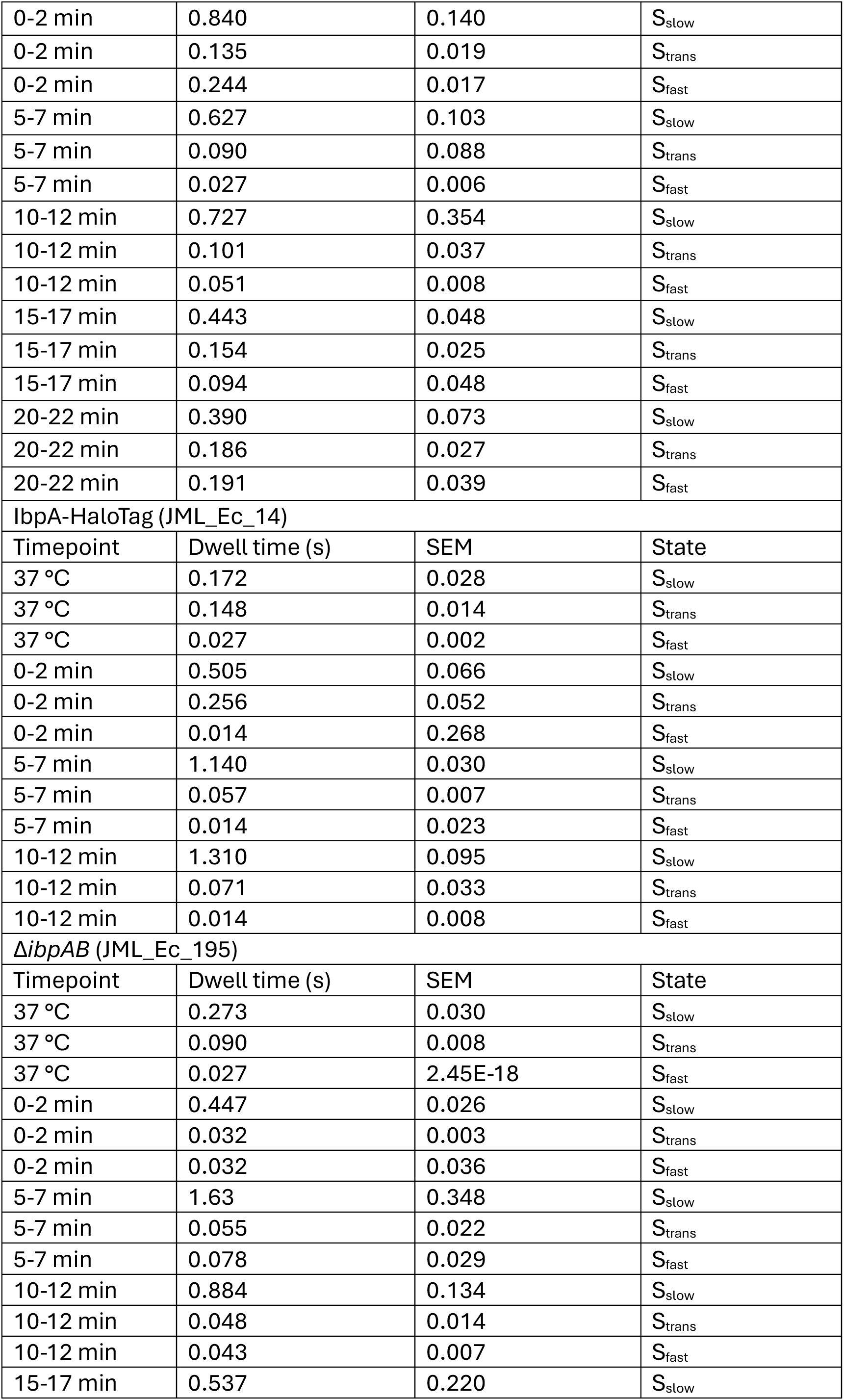
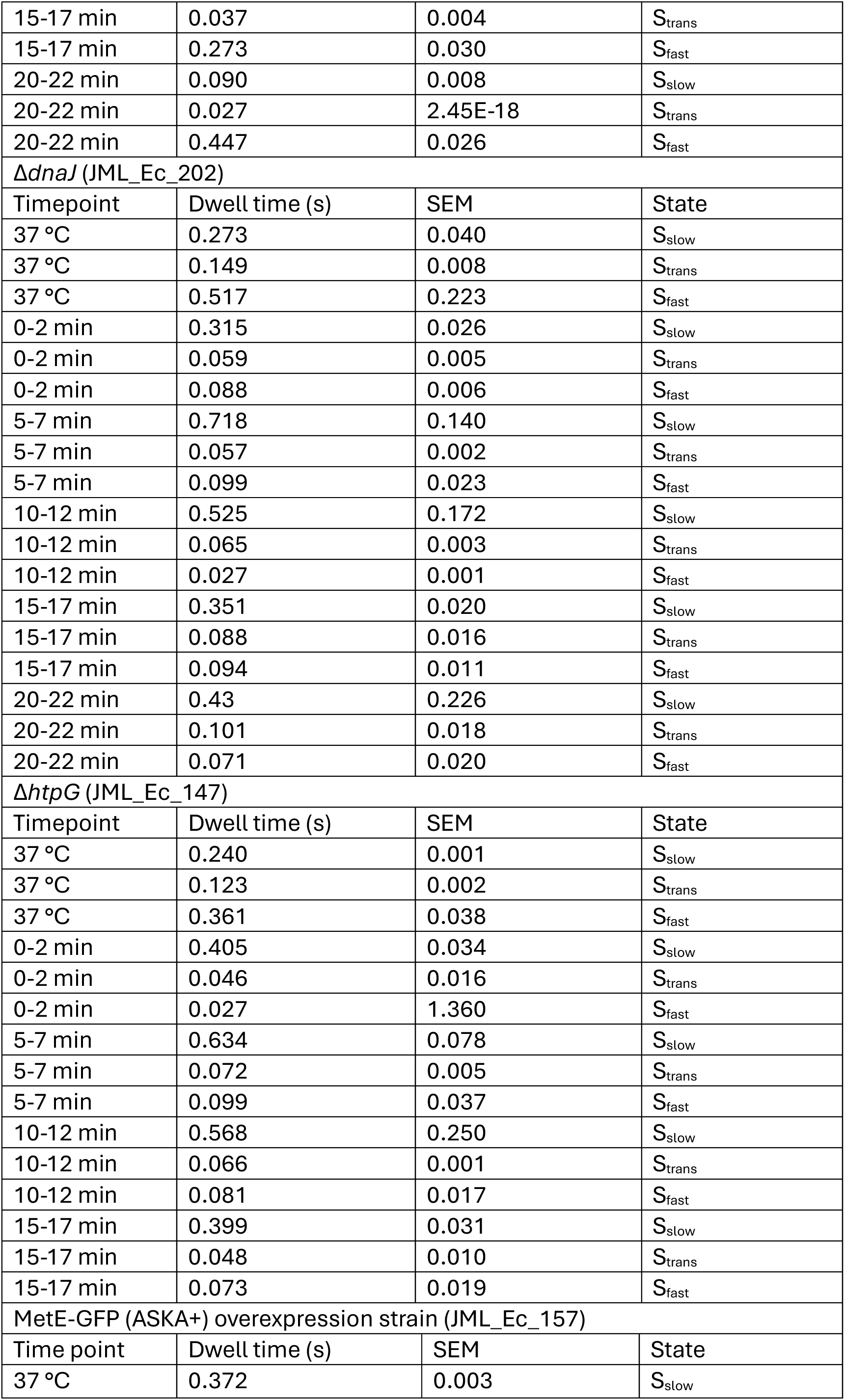
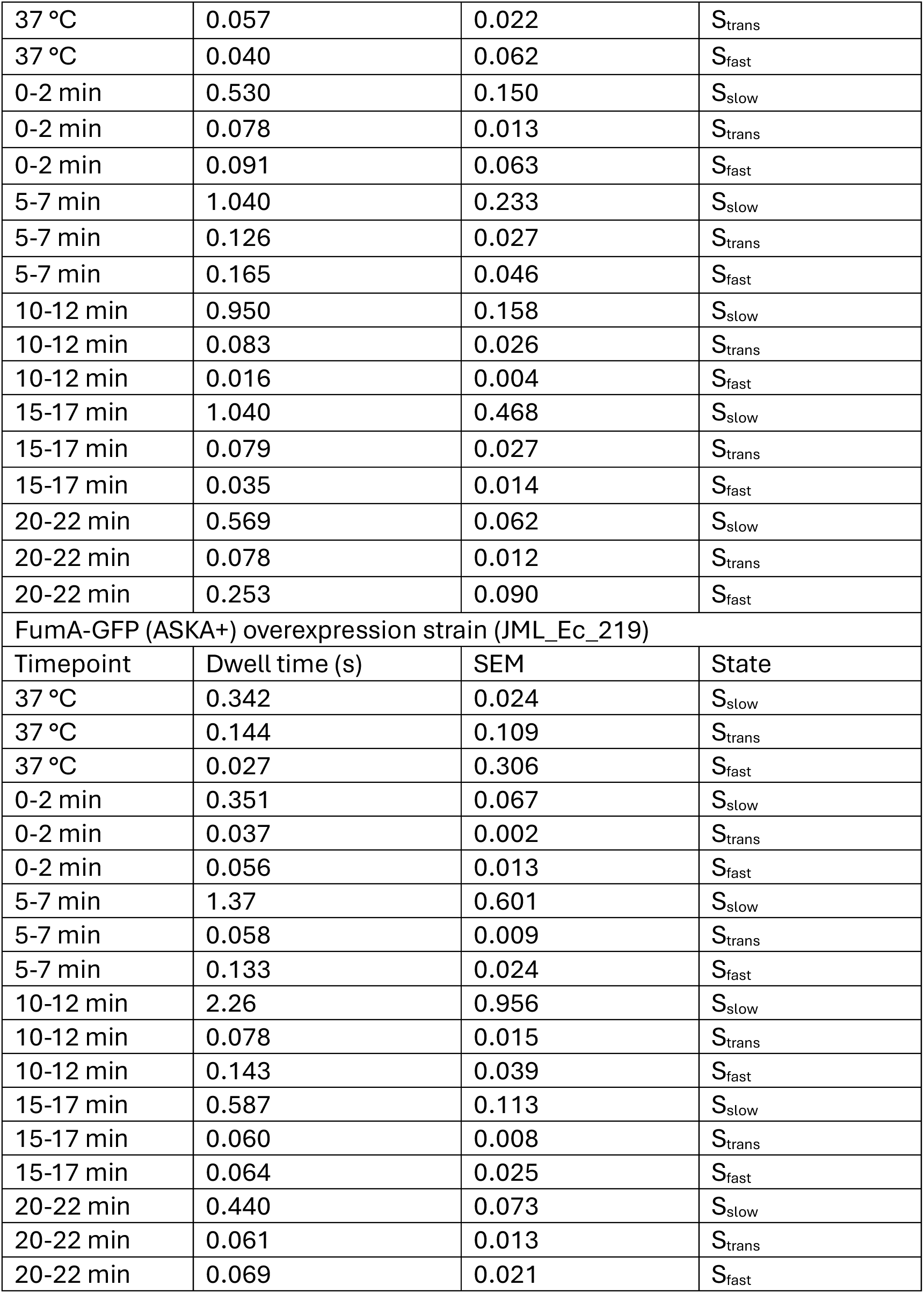
Hidden Markov Model-derived dwell times (±SEM)

**Supplementary Table 4.** Hidden Markov model-derived transition rates. p0, p1, p2 correspond to S_slow,_ S_trans_ and S_fast_ states, respectively.

|  |  |  |  |
| --- | --- | --- | --- |
| DnaK-HaloTag (JML_Ec_05) |  |  |  |
| Transitions | Rate (s<sup>-1</sup>) | Time point | SEM |
| p01 | 4.9 | 37 °C | 0.91 |

| p02 | 0.01 | 37 °C | 2.5E-12 |
| --- | --- | --- | --- |
| p10 | 3.3 | 37 °C | 0.51 |
| p12 | 1.65 | 37 °C | 0.03 |
| p20 | 0.01 | 37 °C | 0.01 |
| p21 | 3.69 | 37 °C | 1.12 |
| p01 | 1.17 | 0-2 min | 0.27 |
| p02 | 0.01 | 0-2 min | 1.13 |
| p10 | 4.99 | 0-2 min | 0.14 |
| p12 | 2.4 | 0-2 min | 1.3E-11 |
| p20 | 0.80 | 0-2 min | 0.42 |
| p21 | 3.3 | 0-2 min | 0.38 |
| p01 | 1.41 | 5-7 min | 0.04 |
| p02 | 0.18 | 5-7 min | 1.87 |
| p10 | 6.26 | 5-7 min | 17.3 |
| p12 | 4.92 | 5-7 min | 0.38 |
| p20 | 0.18 | 5-7 min | 0.78 |
| p21 | 36.7 | 5-7 min | 0.10 |
| p01 | 1.19 | 10-12 min | 0.66 |
| p02 | 0.18 | 10-12 min | 3.60 |
| p10 | 7.25 | 10-12 min | 4.65 |
| p12 | 2.66 | 10-12 min | 0.05 |
| p20 | 4.01 | 10-12 min | 1.09 |
| p21 | 15.7 | 10-12 min | 0.97 |
| p01 | 2.25 | 15-17 min | 0.23 |
| p02 | 0.01 | 15-17 min | 1.21 |
| p10 | 4.59 | 15-17 min | 0.33 |
| p12 | 1.9 | 15-17 min | 1.2E-11 |
| p20 | 0.56 | 15-17 min | 0.35 |
| p21 | 10.1 | 15-17 min | 3.64 |
| p01 | 2.55 | 20-22 min | 0.54 |
| p02 | 0.01 | 20-22 min | 0.70 |
| p10 | 4.3 | 20-22 min | 3.8E-09 |
| p12 | 1.09 | 20-22 min | 4.0E-11 |
| p20 | 0.01 | 20-22 min | 0.14 |
| p21 | 5.22 | 20-22 min | 0.88 |
| IbpA-HaloTag (JML_Ec_14) |  |  |  |
| Transitions | Rate (s<sup>-1</sup>) | Time point | SEM |
| p01 | 5.63 | 37 °C | 0.79 |
| p02 | 0.18 | 37 °C | 0.49 |
| p10 | 5.22 | 37 °C | 0.01 |
| p12 | 1.54 | 37 °C | 0.01 |
| p20 | 0.18 | 37 °C | 0.12 |
| p21 | 36.7 | 37 °C | 0.63 |
| p01 | 1.31 | 0-2 min | 0.50 |
| p02 | 0.67 | 0-2 min | 1.12 |
| p10 | 3.72 | 0-2 min | 14.50 |
| p12 | 0.18 | 0-2 min | 0.13 |
| p20 | 36.7 | 0-2 min | 1.03 |
| p21 | 36.7 | 0-2 min | 2.67 |
| p01 | 0.70 | 5-7 min | 0.40 |
| p02 | 0.18 | 5-7 min | 0.98 |
| p10 | 16.7 | 5-7 min | 2.64 |
| p12 | 0.931 | 5-7 min | 0.93 |
| p20 | 36.7 | 5-7 min | 0.96 |
| p21 | 36.7 | 5-7 min | 0.38 |
| p01 | 0.43 | 10-12 min | 0.10 |
| p02 | 0.34 | 10-12 min | 3.68 |
| p10 | 9.56 | 10-12 min | 1.14 |
| p12 | 4.59 | 10-12 min | 0.09 |
| p20 | 36.7 | 10-12 min | 2.23 |
| p21 | 36.7 | 10-12 min | 5.79 |
| *ΔibpAB* (JML_Ec_195) |  |  |  |
| Transitions | Rate (s<sup>-1</sup>) | Time point | SEM |
| p01 | 3.66 | 37 °C | 0.33 |
| p02 | 0.01 | 37 °C | 1.15 |
| p10 | 7.04 | 37 °C | 0.01 |
| p12 | 1.82 | 37 °C | 1.31E-11 |
| p20 | 0.29 | 37 °C | 0.29 |
| p21 | 1.64 | 37 °C | 0.01 |
| p01 | 2.17 | 0-2 min | 0.12 |
| p02 | 0.11 | 0-2 min | 3.43 |
| p10 | 22.1 | 0-2 min | 0.01 |
| p12 | 4.56 | 0-2 min | 0.04 |
| p20 | 0.01 | 0-2 min | 1.19 |
| p21 | 0.01 | 0-2 min | 3.07 |
| p01 | 0.32 | 5-7 min | 0.06 |
| p02 | 0.30 | 5-7 min | 1.41 |
| p10 | 9.43 | 5-7 min | 4.33 |
| p12 | 8.87 | 5-7 min | 0.13 |
| p20 | 2.53 | 5-7 min | 8.49 |
| p21 | 10.3 | 5-7 min | 29.6 |
| p01 | 1.1 | 10-12 min | 0.27 |
| p02 | 0.0 | 10-12 min | 4.31 |
| p10 | 12.4 | 10-12 min | 6.69 |
| p12 | 8.57 | 10-12 min | 0.32 |
| p20 | 0.01 | 10-12 min | 0.32 |
| p21 | 23.1 | 10-12 min | 14.3 |
| p01 | 1.64 | 15-17 min | 0.38 |
| p02 | 0.22 | 15-17 min | 1.80 |
| p10 | 14.3 | 15-17 min | 0.59 |
| p12 | 12.9 | 15-17 min | 0.10 |
| p20 | 1.13 | 15-17 min | 2.15 |
| p21 | 17.6 | 15-17 min | 2.70 |
| p01 | 1.44 | 20-22 min | 0.60 |
| p02 | 0.17 | 20-22 min | 2.47 |
| p10 | 17.9 | 20-22 min | 0.82 |
| p12 | 2.48 | 20-22 min | 0.02 |
| p20 | 0.53 | 20-22 min | 0.28 |
| p21 | 3.11 | 20-22 min | 1.55 |
| *ΔdnaJ* (JML_Ec_202) |  |  |  |
| Transitions | Rate (s<sup>-1</sup>) | Time point | SEM |
| p01 | 3.65 | 37 °C | 0.39 |
| p02 | 0.01 | 37 °C | 0.31 |
| p10 | 5.44 | 37 °C | 4.5E-11 |
| p12 | 1.29 | 37 °C | 1.4E-12 |
| p20 | 0.01 | 37 °C | 0.32 |
| p21 | 1.92 | 37 °C | 0.59 |
| p01 | 2.99 | 0-2 min | 0.33 |
| p02 | 0.19 | 0-2 min | 1.42 |
| p10 | 10.1 | 0-2 min | 1.30 |
| p12 | 7.01 | 0-2 min | 0.61 |
| p20 | 0.017 | 0-2 min | 0.36 |
| p21 | 11.4 | 0-2 min | 0.59 |
| p01 | 1.13 | 5-7 min | 0.21 |
| p02 | 0.26 | 5-7 min | 0.78 |
| p10 | 7.08 | 5-7 min | 0.15 |
| p12 | 10.5 | 5-7 min | 0.08 |
| p20 | 1.87 | 5-7 min | 0.11 |
| p21 | 8.23 | 5-7 min | 2.21 |
| p01 | 1.73 | 10-12 min | 0.94 |
| p02 | 0.18 | 10-12 min | 2.54 |
| p10 | 7.06 | 10-12 min | 58.8 |
| p12 | 8.42 | 10-12 min | 0.06 |
| p20 | 0.01 | 10-12 min | 1.02 |
| p21 | 36.7 | 10-12 min | 1.39 |
| p01 | 2.69 | 15-17 min | 0.99 |
| p02 | 0.17 | 15-17 min | 5.12 |
| p10 | 5.69 | 15-17 min | 2.98 |
| p12 | 5.66 | 15-17 min | 0.76 |
| p20 | 0.01 | 15-17 min | 2.16 |
| p21 | 10.6 | 15-17 min | 0.73 |
| p01 | 2.32 | 20-22 min | 1.15 |
| p02 | 0.01 | 20-22 min | 0.70 |
| p10 | 4.69 | 20-22 min | 60.9 |
| p12 | 5.25 | 20-22 min | 0.17 |
| p20 | 1.11 | 20-22 min | 2.09 |
| p21 | 13 | 20-22 min | 15.9 |
| *ΔhtpG* (JML_Ec_147) |  |  |  |
| Transitions | Rate (s<sup>-1</sup>) | Time point | SEM |
| p01 | 0.01 | 37 °C | 0.03 |
| p02 | 7 | 37 °C | 0.16 |
| p10 | 1.11 | 37 °C | 3.3E-10 |
| p12 | 0.01 | 37 °C | 1.7E-10 |
| p20 | 2.76 | 37 °C | 0.02 |
| p21 | 2.47 | 37 °C | 0.18 |
| p01 | 0.01 | 0-2 min | 0.21 |
| p02 | 13.4 | 0-2 min | 4.34 |
| p10 | 3.44 | 0-2 min | 0.43 |
| p12 | 0.85 | 0-2 min | 8.4E-08 |
| p20 | 0.01 | 0-2 min | 0.90 |
| p21 | 1.32 | 0-2 min | 9.15 |
| p01 | 0.26 | 5-7 min | 0.26 |
| p02 | 10.2 | 5-7 min | 1.01 |
| p10 | 3.7 | 5-7 min | 2.74 |
| p12 | 6.01 | 5-7 min | 0.12 |
| p20 | 4.13 | 5-7 min | 0.33 |
| p21 | 1.26 | 5-7 min | 2.71 |
| p01 | 0.50 | 10-12 min | 0.37 |
| p02 | 6.01 | 10-12 min | 2.13 |
| p10 | 9.12 | 10-12 min | 1.08 |
| p12 | 3.72 | 10-12 min | 0.12 |
| p20 | 8.59 | 10-12 min | 1.89 |
| p21 | 2.36 | 10-12 min | 4.34 |
| p01 | 0.15 | 15-17 min | 0.25 |
| p02 | 9 | 15-17 min | 1.37 |
| p10 | 11.9 | 15-17 min | 0.45 |
| p12 | 0.90 | 15-17 min | 0.06 |
| p20 | 12.9 | 15-17 min | 4.85 |
| p21 | 0.01 | 15-17 min | 3.47 |
| MetE-GFP (ASKA+) overexpression strain (JML_Ec_157) |  |  |  |
| Transitions | Rate (s<sup>-1</sup>) | Time point | SEM |
| p01 | 2.54 | 37 °C | 0.06 |
| p02 | 0.14 | 37 °C | 0.5 |
| p10 | 3.45 | 37 °C | 0.10 |
| p12 | 14 | 37 °C | 0.01 |
| p20 | 0.14 | 37 °C | 0.20 |
| p21 | 24.6 | 37 °C | 1.18 |
| p01 | 1.88 | 0-2 min | 0.30 |
| p02 | 0.01 | 0-2 min | 0.59 |
| p10 | 8.98 | 0-2 min | 1.27 |
| p12 | 3.8 | 0-2 min | 0.02 |
| p20 | 0.24 | 0-2 min | 1.56 |
| p21 | 10.8 | 0-2 min | 5.69 |
| p01 | 0.85 | 5-7 min | 0.27 |
| p02 | 0.12 | 5-7 min | 2.51 |
| p10 | 6.06 | 5-7 min | 5.7E-09 |
| p12 | 1.85 | 5-7 min | 0.04 |
| p20 | 0.01 | 5-7 min | 1.09 |
| p21 | 6.07 | 5-7 min | 3.01 |
| p01 | 3.11 | 10-12 min | 0.25 |
| p02 | 0.32 | 10-12 min | 2.28 |
| p10 | 14.6 | 10-12 min | 2.44 |
| p12 | 7.67 | 10-12 min | 0.19 |
| p20 | 9.08 | 10-12 min | 3.96 |
| p21 | 9.54 | 10-12 min | 4.38 |
| p01 | 0.42 | 15-17 min | 0.11 |
| p02 | 0.54 | 15-17 min | 1.50 |
| p10 | 5.07 | 15-17 min | 6.83 |
| p12 | 7.53 | 15-17 min | 0.11 |
| p20 | 5.93 | 15-17 min | 4.97 |
| p21 | 22.5 | 15-17 min | 24.8 |
| p01 | 1.41 | 20-22 min | 0.06 |
| p02 | 0.35 | 20-22 min | 0.94 |
| p10 | 8.3 | 20-22 min | 0.71 |
| p12 | 4.45 | 20-22 min | 0.07 |
| p20 | 1.31 | 20-22 min | 1.32 |
| p21 | 2.65 | 20-22 min | 1.49 |
| FumA-GFP (ASKA+) overexpression strain (JML_Ec_219) |  |  |  |
| Transitions | Rate (s<sup>-1</sup>) | Time point | SEM |
| p01 | 2.74 | 37 °C | 0.56 |
| p02 | 0.18 | 37 °C | 1.08 |
| p10 | 4.68 | 37 °C | 61.2 |
| p12 | 2.27 | 37 °C | 0.38 |
| p20 | 0.18 | 37 °C | 0.24 |
| p21 | 36.7 | 37 °C | 12.2 |
| p01 | 2.81 | 0-2 min | 0.41 |
| p02 | 0.04 | 0-2 min | 0.80 |
| p10 | 14.2 | 0-2 min | 0.57 |
| p12 | 12.5 | 0-2 min | 0.02 |
| p20 | 0.20 | 0-2 min | 0.68 |
| p21 | 17.7 | 0-2 min | 5.14 |
| p01 | 0.68 | 5-7 min | 0.16 |
| p02 | 0.05 | 5-7 min | 3.92 |
| p10 | 13.6 | 5-7 min | 1.21 |
| p12 | 3.8 | 5-7 min | 0.06 |
| p20 | 3.01 | 5-7 min | 0.90 |
| p21 | 4.49 | 5-7 min | 1.86 |
| p01 | 0.40 | 10-12 min | 0.48 |
| p02 | 0.04 | 10-12 min | 4.25 |
| p10 | 10.2 | 10-12 min | 2.32 |
| p12 | 2.61 | 10-12 min | 0.02 |
| p20 | 2.72 | 10-12 min | 0.51 |
| p21 | 4.26 | 10-12 min | 1.66 |
| p01 | 1.56 | 15-17 min | 0.27 |
| p02 | 0.14 | 15-17 min | 6.89 |
| p10 | 9.74 | 15-17 min | 1.16 |
| p12 | 6.92 | 15-17 min | 0.02 |
| p20 | 3.19 | 15-17 min | 2.85 |
| p21 | 12.5 | 15-17 min | 6.91 |
| p01 | 2.27 | 20-22 min | 0.29 |
| p02 | 0.01 | 20-22 min | 3.35 |
| p10 | 11.2 | 20-22 min | 0.36 |
| p12 | 5.15 | 20-22 min | 0.01 |
| p20 | 0.34 | 20-22 min | 0.43 |
| p21 | 14.2 | 20-22 min | 2.77 |

**Supplementary Table 5.** Strains used in this study.

| Identifier | Difference from background | Source or reference |
| --- | --- | --- |
| MG1655 | *F- lambda- ilvG- rfb- 50 rph-1* | <sup>63</sup> |
| JML_Ec_05 | *dnaK::dnaK-HaloTag-frt-kan-frt* | The HaloTag and kanamycin cassette were inserted into the C-terminus of the native chromosomal *dnaK* gene through λ-RED recombination using the primer pair JMP_Ec_01_Fw and JMP_Ec_02_Rv. The construct was then transferred to MG1655 using P1 phage transduction. |
| JML_Ec_14 | *ibpA::ibpA-HaloTag-frt-kan-frt* | The HaloTag and kanamycin cassette were inserted into the C-terminus of the native chromosomal *ibpA* gene through λ-RED recombination using the primer pair JMP_Ec_38_Fw and JMP_Ec_39_Rv. The construct was then transferred to MG1655 using P1 phage transduction. |
| JML_Ec_20 | *dnaJ::dnaJ-HaloTag-frt-kan-frt* | The HaloTag and kanamycin cassette were inserted into the C- |
| | | terminus of the native chromosomal *dnaJ* gene through λ-RED recombination using the primer pair JMP_Ec_34_Fw and JMP_Ec_35_Rv. The construct was then transferred to MG1655 using P1 phage transduction. |
| JML_Ec_29 | *dnaK::dnaK-HaloTag-frt* | The kanamycin resistance cassette was excised from JML_Ec_05 by expressing the Flp site-specific recombinase from pCP20. |
| JML_Ec_136 | *metE::metE-HaloTag-frt-kan-frt* | The HaloTag and kanamycin cassette were inserted into the C-terminus of the native chromosomal *metE* gene through λ-RED recombination using primer pair JMP_Ec_350_Fw and JMP_Ec_351_Rv. |
| JML_Ec_141 | *glcB::glcB-HaloTag-frt-kan-frt* | The HaloTag and kanamycin cassette were inserted into the C-terminus of the native chromosomal *glcB* gene through λ-RED recombination using the primer pair JMP_Ec_354_Fw and JMP_Ec_355_Rv. |
| JML_Ec_147 | *dnaK::dnaK-HaloTag-frt, ΔhtpG::frt-kan-frt* | The *htpG* deletion (kanamycin insertion) was transferred from the Escherichia coli single-gene deletion Keio collection strain JW0462-1<sup>64</sup> to JML_Ec_29 through P1 transduction. |
| JML_Ec_157 | *dnak::dnak-HaloTag-frt, pCA24N-metE-gfp* | Transformation of pCA24N-*metE-gfp* from the ASKA+ library strain JW3805<sup>42</sup> into JML_Ec_29. |
| JML_Ec_164 | *dnak:dnak<sup>S427P</sup>-HaloTag-frt-kan-frt* | The DnaK<sup>S427P</sup>-HaloTag was generated using Gibson assembly and site-directed mutagenesis. First, the *dnaK* and HaloTag-kanamycin cassette was cloned into a pUC19 backbone using the following primer pairs: JMP_Ec_140_Fw/ JMP_Ec_141_Rv and JMP_Ec_142_Fw/ JMP_Ec_143_Rv, respectively. The S427P single-site mutation was generated using the overlapping |
| | | primer pair JMP_516_Fw and JMP_517_Rv. The full DnaK<sup>S427P</sup>-HaloTag construct was then amplified using the primer pair JMP_Ec_252_Fw/JMP_Ec_02_Rv and inserted to the native DnaK locus using λ-RED recombination. |
| JML_Ec_194 | *dnak::dnaK<sup>V436F</sup>-HaloTag-frt-kan-frt* | The DnaK<sup>V436F</sup>-HaloTag construct was generated by Gibson assembly. The DnaK<sup>V436F</sup> fragment was amplified from pNB64 (JMP_Ec_374_Fw/JMP_Ec_375_Rv) and fused to HaloTag-kanamycin cassette (JMP_Ec_376_Fw/JMP_Ec_377_Rv) using Gibson assembly. The pUC19 backbone was amplified using JMP_Ec_378_Fw/ JMP_Ec_379_Rv. The full DnaK<sup>V436F</sup>-HaloTag construct was then amplified using JMP_Ec_254_Fw/ JMP_Ec_02_Rv from this cloning vector and inserted into the native chromosomal *dnak* locus through λ-RED recombination. |
| JML_Ec_195 | *ΔibpAB::frt-kan-frt, dnak::dnaK-HaloTag-frt* | The *ibpAB* operon was replaced by a kanamycin cassette through λ-RED recombination using the primer pair JMP_Ec_396_Fw/ JMP_Ec_397_Rv. The construct was then transferred to JML_Ec_29 cells using P1 phage transduction. |
| JML_Ec_202 | *ΔdnaJ::frt-kan-frt, dnak::dnaK-HaloTag-frt* | The *dnaJ* gene was replaced by a kanamycin cassette through λ-RED recombination using the primer pair JMP_340_Fw/ JMP_341_Rv. The construct was then transferred to JML_Ec_29 cells through another round of λ-RED recombination using the primer pair JMP_Ec_394_Fw/ JMP_Ec_395_Rv. |
| JML_Ec_219 | *dnak::dnak-HaloTag-frt, pCA24N-fumA-gfp* | Transformation of pCA24N-fumA-gfp from the ASKA+ library strain JW1604<sup>42</sup> into JML_Ec_29. |
| JML_Ec_249 | *metE::metE-gfp-cm* | The *metE-gfp* construct was amplified from the ASKA+ collection strain JW3805<sup>42</sup> and inserted into the *metE* chromosomal locus by λ-RED recombination using the primer pair JMP_Ec_573_FW+ JMP_Ec_574_RV |
| JML_Ec_250 | *fumA::fumA-gfp-cm* | The *fumA-gfp* construct was amplified from the ASKA+ collection strain JW1604<sup>42</sup> and inserted into the *fumA* chromosomal locus by λ-RED recombination using the primer pair JMP_Ec_569_FW/ JMP_Ec_570_Rv. |

**Supplementary Table 6.**
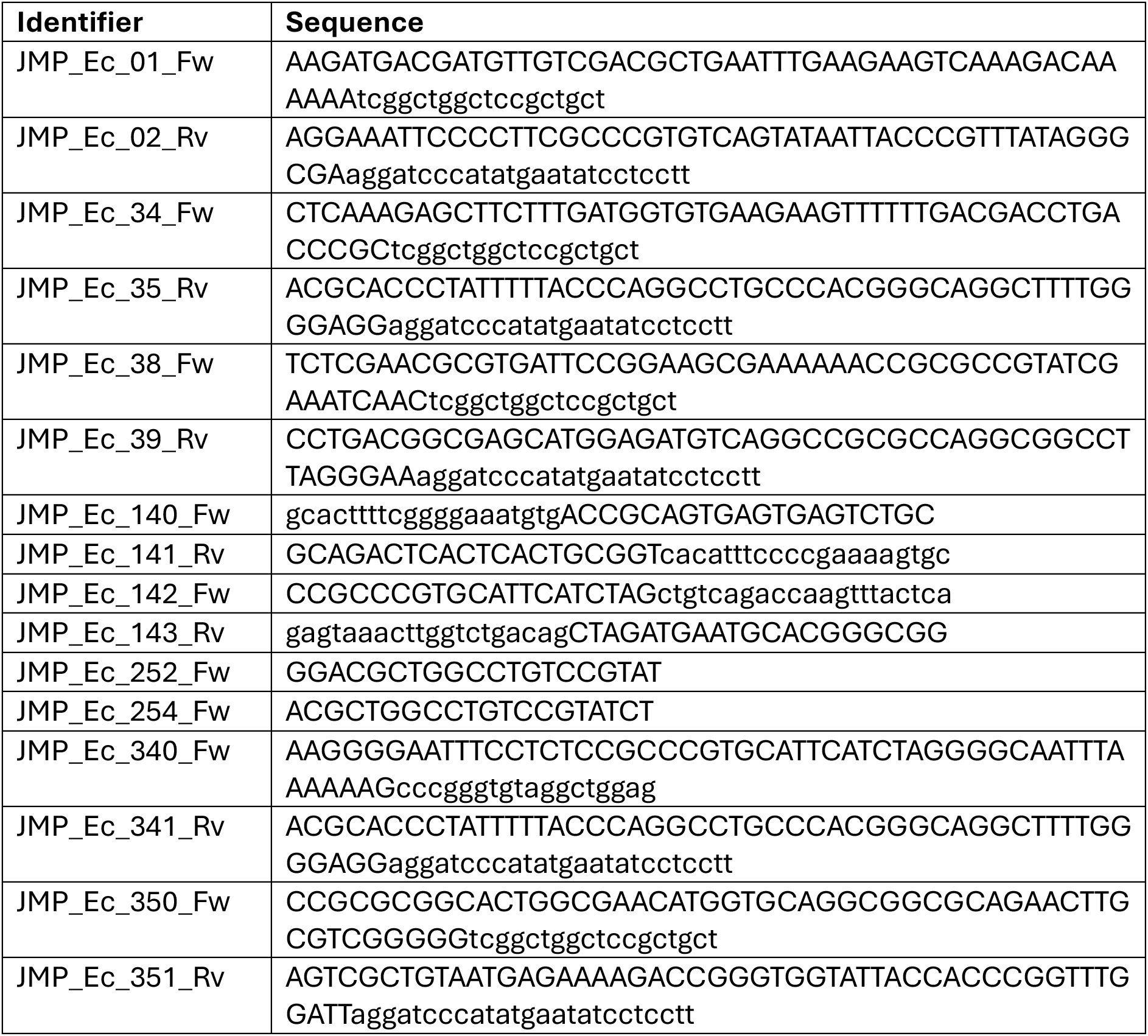
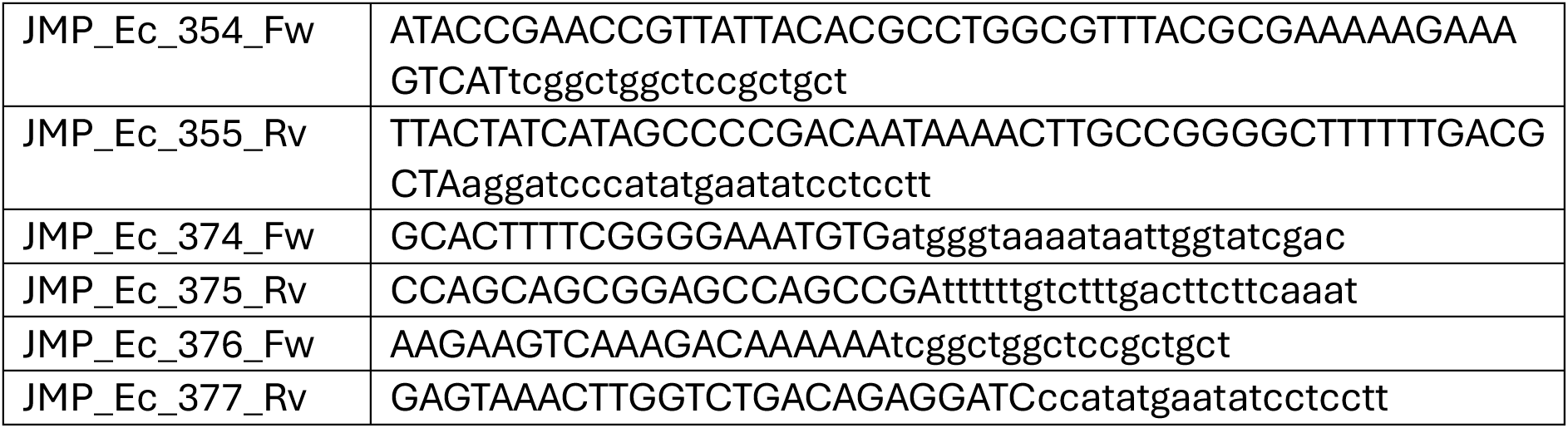
Oligonucleotides used in this study.

**Supplementary Table 7.**
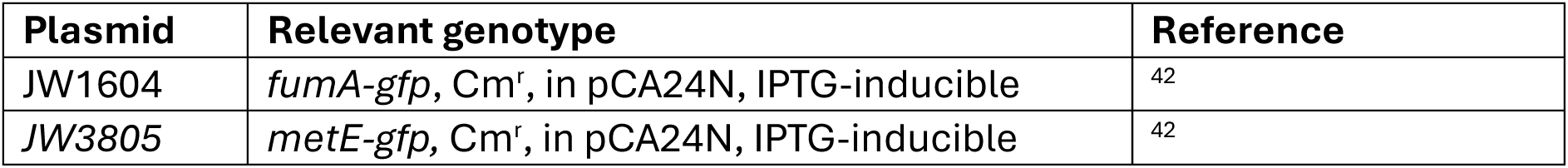
Plasmids used in this study.

